# BBrowser: Making single-cell data easily accessible

**DOI:** 10.1101/2020.12.11.414136

**Authors:** Tri Le, Tan Phan, Minh Pham, Dat Tran, Loc Lam, Tung Nguyen, Thao Truong, Hy Vuong, Tam Luu, Nam Phung, Ngan Pham, Trang Nguyen, Oanh Pham, An Nguyen, Huy Nguyen, Hao Tran, Loc Tran, Ha An Nguyen, Thanh Tran, Nhung Nguyen, Ngoc Tran, Cecilie Boysen, Uyen Nguyen, Vy Pham, Theodore Kim, Ngoc Pham, Tristan Gill, Son Pham

## Abstract

BioTuring’s BBrowser is a software solution that helps scientists effectively analyze single-cell omics data. It combines big data with big computation and modern data visualization to create a unique platform where scientists can interact and obtain important biological insights from the massive amounts of single-cell data. BBrowser has three main components: a curated single-cell database, a big-data analytics layer, and a data visualization module. BBrowser is available for download at: https://bioturing.com/bbrowser/download.

## INTRODUCTION

The emergence of single-cell sequencing technology in recent years has raised numerous challenges, starting with large scale computationally intensive data analytics, to data curation, handling, and comparisons, onto visualization. Before the single-cell era, each data set typically contained up to hundreds of data points, now hundreds of thousands of single cells are each representing a data point in thousands of dimensions in different multi-omics modalities. Besides the number of data points for each experiment, the number of data sets also increases exponentially. This raises a huge challenge to collect, curate, and store all these data in a unified structure. Currently, the total number of single cells analyzed, here considering only published papers including scRNAseq data, is more than several dozens of millions of cells, each in at least 20,000 dimensions. This yields an urgent need for a new set of effective algorithms to analyze the data. Additionally, given the high number of data points in high dimensional space, a completely new suite of visualization packages need to be designed to allow scientists to interact with the data. We present BBrowser to address these challenges.

## A CURATED SINGLE-CELL DATABASE

By the time of writing this paper, the Pubmed database has 465 papers with single-cell data. BioTuring has curated 305 data sets from here and created a database with more than 15 million cells including 376 cell types from diverse organs and 60 diseases (Figure 1). For each data set, we collect metadata, expression matrices, low-dimensional embedding coordinates (e.g., t-SNE, UMAP), and clustering information. Importantly, we try to reproduce some key findings in the papers. If we fail to reproduce the finding in the paper, we contact the authors for clarification.

**Figure 1.**
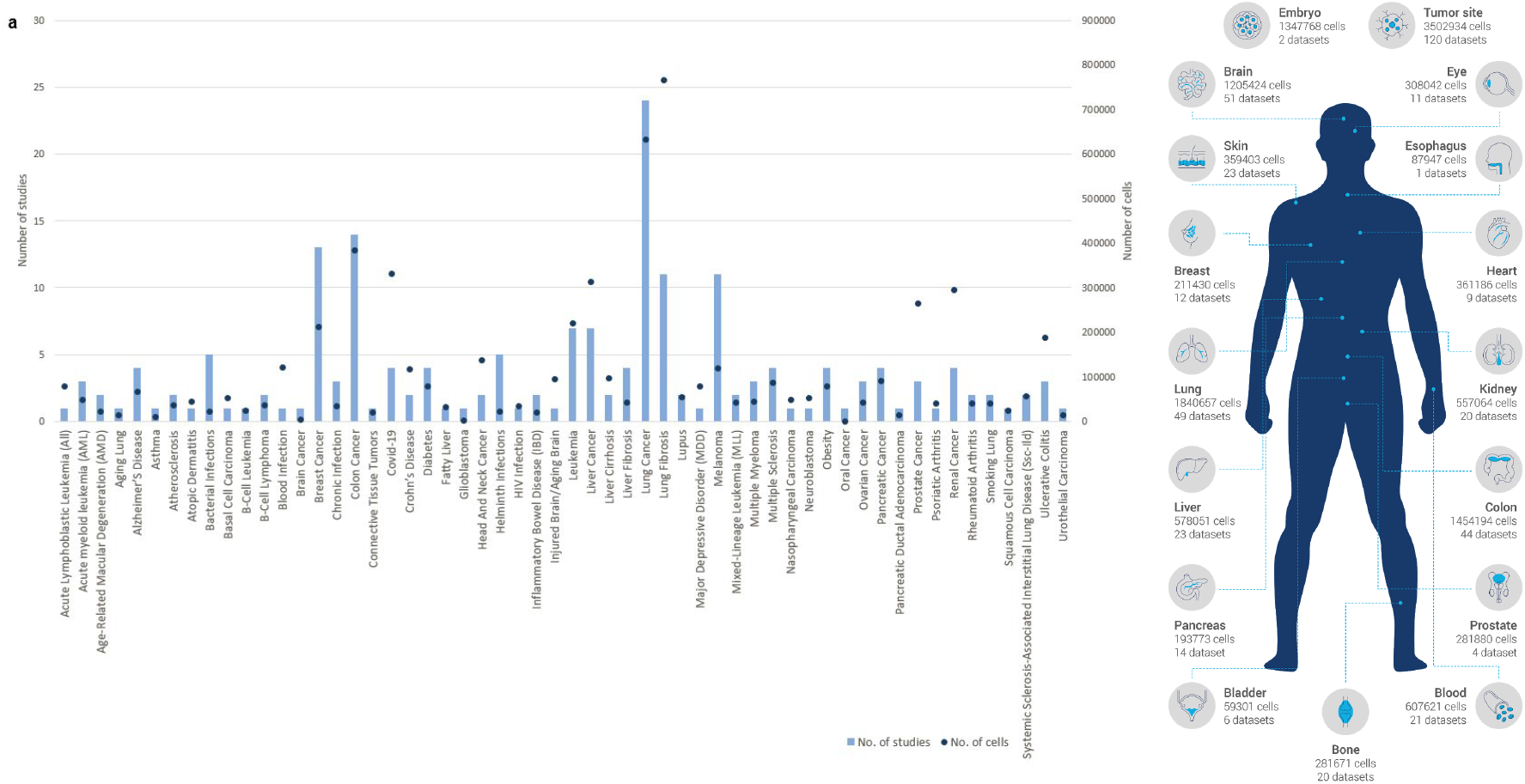
An overview of BioTuring Curated Single-cell Database with 15 million cells. (a) Number of cells and studies across 56 diseases. (b) Number of cells and studies in 17 tissues.

A key step to make data comparable between studies is the use of a controlled vocabulary. We built our own controlled vocabulary and ontology for cell types, diseases, age, drug, treatment, and anatomy. Our database uses the Cell Line ontology (Sarntivijai et al., 2014), Disease Ontology (Schriml et al. 2012), and MeSH (Lipscomb, 2000) as references for our controlled vocabularies. To avoid the biases in cell type annotation from different authors/laboratories, we also applied our own cell type prediction algorithm across all data sets.

With the huge curated single-cell database in place and constantly expanding, the question is how to utilize this database. A straightforward way is to allow users to download and analyze each dataset individually, which BBrowser allows you to do. Hower, here we show that by applying an advanced computational layer on top of this database, one can begin to ask complicated queries and summarize results across the entire database. In the next section, we describe several important algorithms in the advanced computational layer, including *Cell Search* and *Talk2Data*.

## BIG COMPUTATION ON BIG DATA

### Cell Search

One of the most common tasks in single-cell analysis involves cell type annotation. Often, for a selected cell population resulting from a chosen clustering method, scientists need to manually check the expression of these cells against a list of marker genes from literature for each cell type and choose the cell type that best matches with the list. With the curated and well-annotated population of cells in the BioTuring database, we can formulate this task as finding the populations in the database that have gene expression profiles that best match the selected population, and transfer the annotation(s) from the matched populations to the query one (Figure 2). This is the key idea of Cell Search. Below, we describe the Cell Search algorithm.

**Figure 2.**
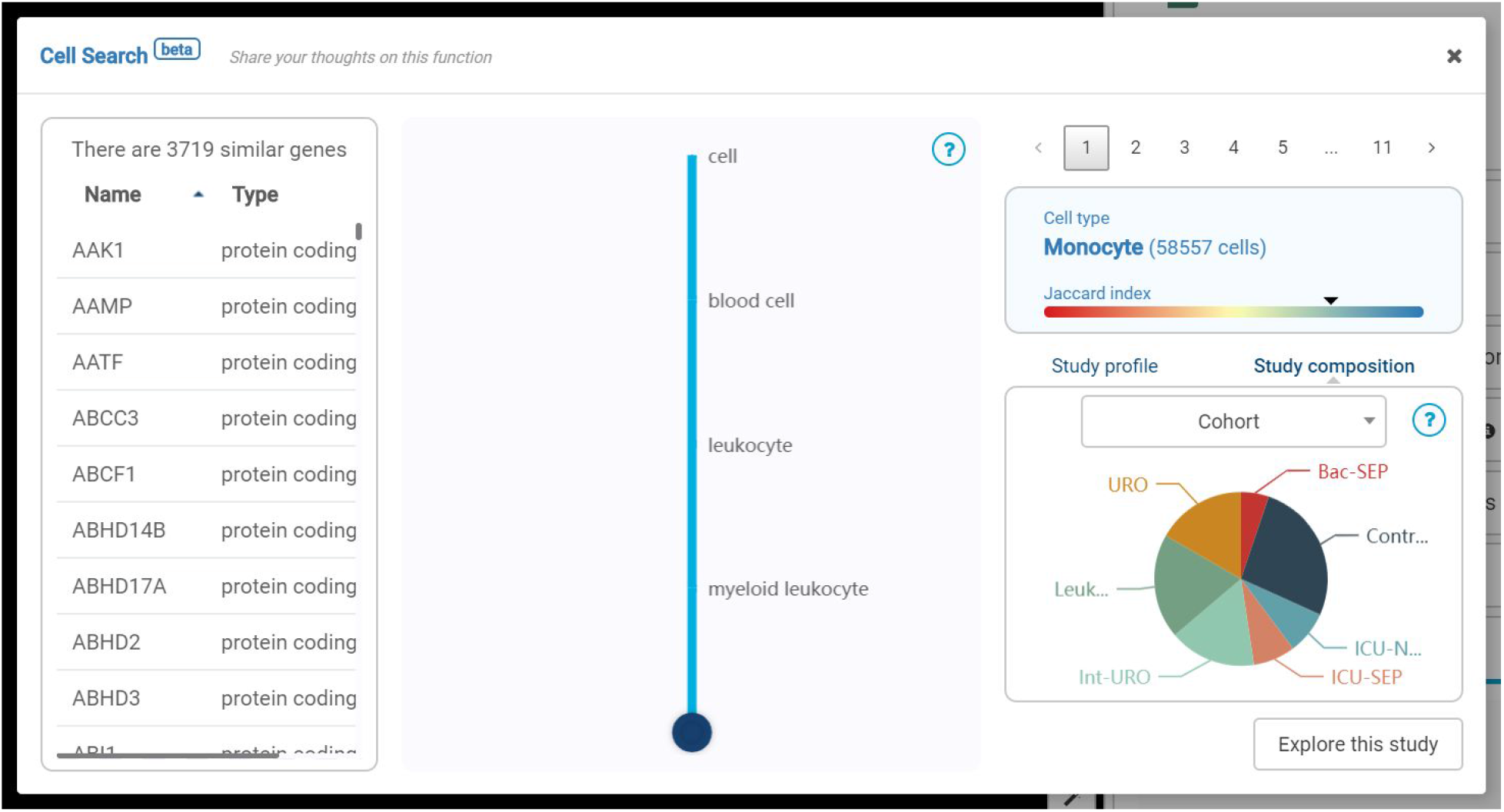
The interface of a cell search result in BBrowser. In the middle, there is a cell tree summarizing all the cell populations that matches the cells of query. On the right are the studies that contain a corresponding group of cells. On the left is the list of genes that are shared with the cells of the query.

The Cell Search algorithm has two parts: *indexing* and *searching*. For each cluster (graph-based clustering) in the database, we create a profile, each is a vector in 20,000 dimensional gene space. When a user selects a population of cells to search, we generate a corresponding profile of that population and perform a query to find similar profiles in the index database.

### Talk2Data

The use of controlled vocabularies in BioTuring Database allows us to perform many analytical queries and filtering across different data sets. For example, a user who wants to look for T cells can use the *cell type filter* in the software. This filter will then use the cell type metadata, which has already been standardized into a controlled label in an ontology, to look for T cells and even sub-types of T cells.

The use of controlled vocabularies across datasets enables us to create a big single data set with dozens of millions of cells. This huge table helps reveal many biological insights that consistently appear in different datasets (from different laboratories) and emerges only with a large number of data points. Users can interact with this database in BBrowser via the Talk2Data interface (Figure 3). Below are some questions that Talk2Data can help answer:

- *What genes are co-expressed with FOXP3 in regulatory T cells in lung cancer?*
- *What is the list of gene markers of regulatory T cells in cancer?*

**Figure 3.**
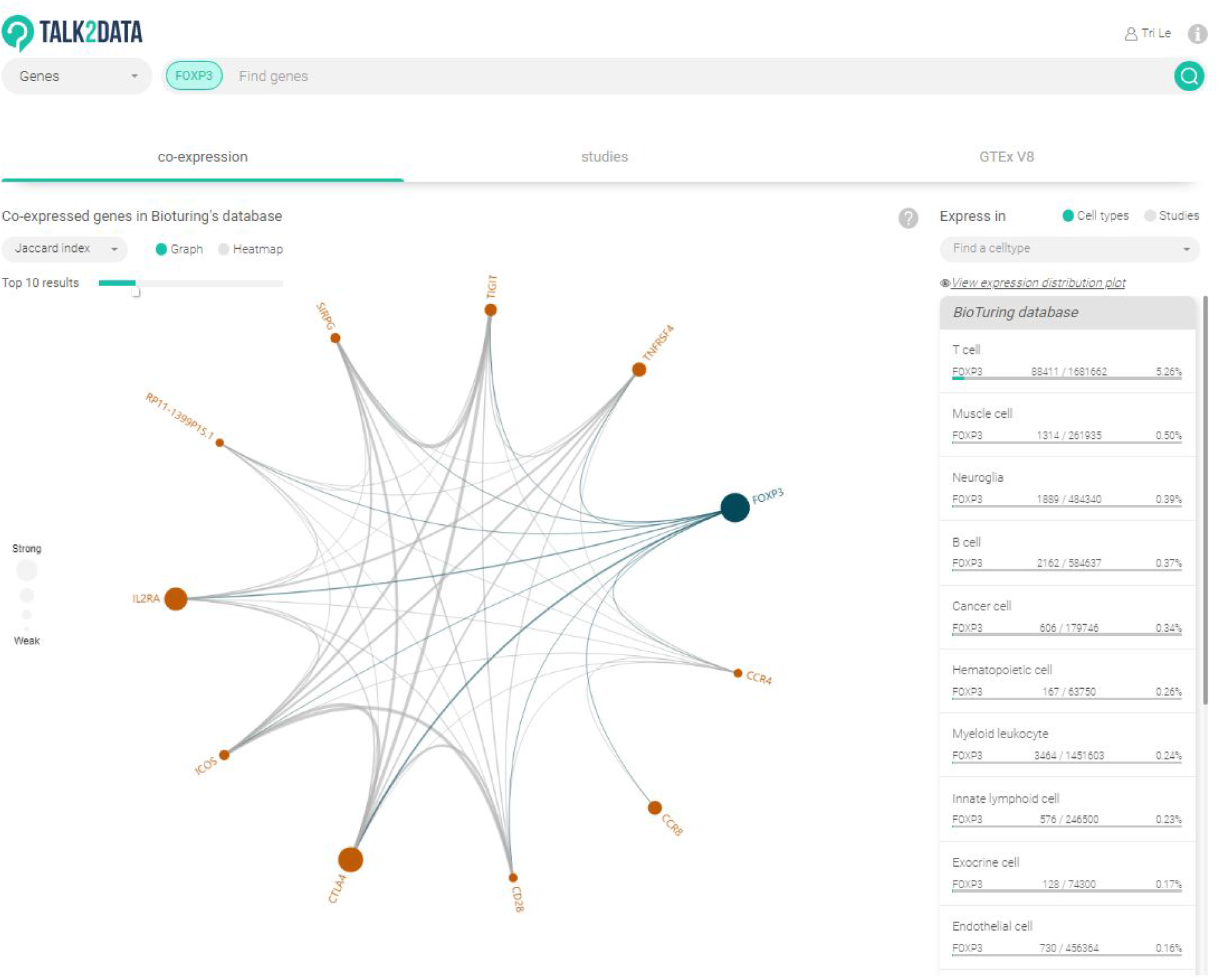
Interface of Talk2Data for gene-gene correlation network. Size of points indicates the level of correlation.

## MODERN DATA VISUALIZATION FOR END USERS

### Access the Database

To access curated BioTuring Single-cell Database, anyone can go to our website (https://www.bioturing.com/bbrowser) and download BBrowser. This is a desktop application and needs internet connection to launch. The default interface of BBrowser is a library of single-cell publications (Figure 4). This interface shows basic information about each paper as well as its single-cell data set such as the number of cells, species, sequencing technology, and the type of omics. We developed a text search engine that helps a user perform standard keyword search and apply various filtering options. To access a particular data set, the user just needs to click “Download” and then “Explore” it. BBrowser will download necessary data to the local computer and use local computing resources for visualizations and analysis.

**Figure 4.**
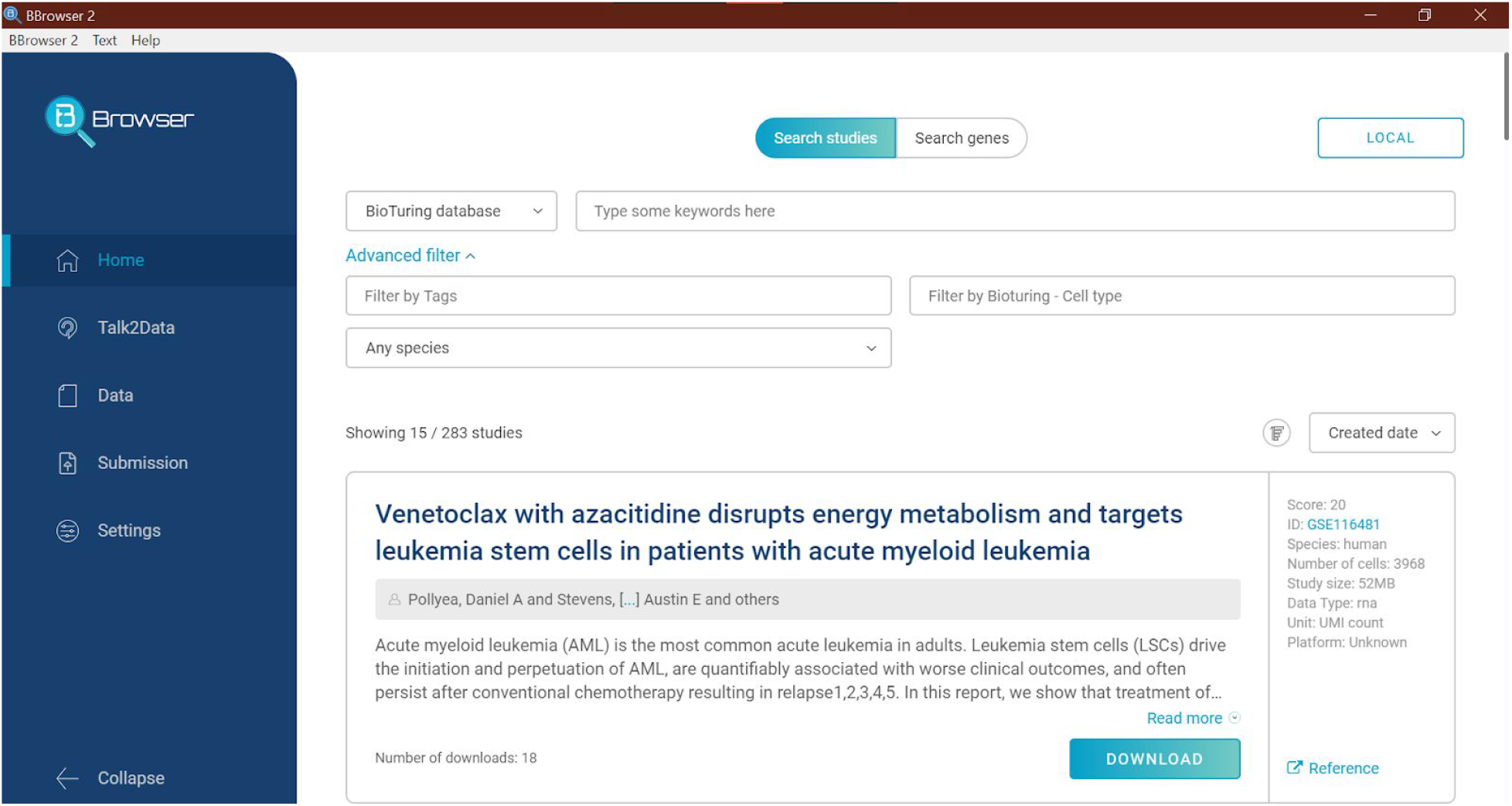
The default interface of BBrowser. This is an interface to query our Database of single-cell data sets.

### General Visualization

When a user accesses a data set in BBrowser, the software will use a scatter plot as a primary visualization (Figure 5). This is called the Main Plot, where each data point is a single cell (or a single spot for spatial data), coordinates are the embeddings such as t-SNE or UMAP (or spatial coordinates). By default, the color of cells is the louvain clustering result. The user can query one or more genes to change the color into expression values. The Main Plot is interactive, which means a user can zoom, drag, rotate (in 3D coordinates), filter, and select cells. Users can select a population of cells by using lasso selection, groups in metadata, or expression of one or more genes. We reengineered a javascript visualization library to be able to visualize millions of cells on standard laptops. To the best of our knowledge, there is no other tool that can achieve an interactive visualization on that high number of cells.

**Figure 5.**
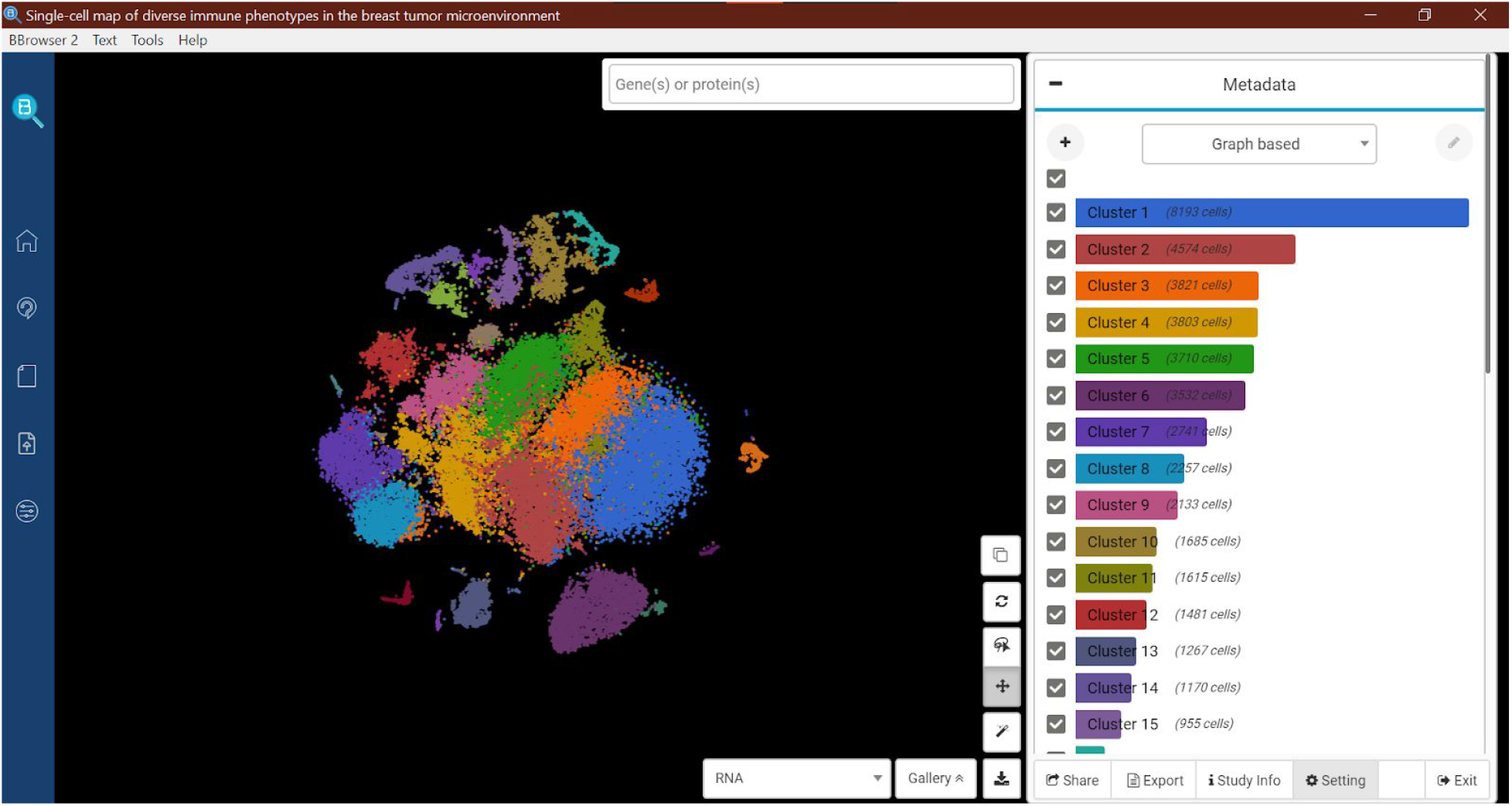
The interface of BBrowser for a single-cell data set of breast cancer (Azizi et al., 2018). In this study, the Main Plot is the t-SNE of 47,016 cells. Cells are colored with the graph-based clustering result from the mRNA expression.

### Filter and Selection

An important operation in single-cell analysis is the ability to correctly and consistently choose a population of cells. In order to assist the robustness of selection, BBrowser has a special function to filter cells using a set of criteria. The most basic filter is using the checkboxes in the Metadata panel (Figure 6), where the user can simply uncheck the cells that are not of interest. Any selection afterwards will only apply on the visible cells in the Main Plot, which is easier to reproduce. Users may often need to perform selections with complex criteria. For example, the user wants to select the CD3D+ CD4+ cells from the tumor samples of patient BC1 to BC3. In this case, the user can create such criteria in the Filter panel (Figure 7). We designed a user-friendly interface to help users create complex combinations of logical conditions using metadata and expression. Such combinations are automatically saved and can be reused for other purposes. With this flexibility to filter and select cells, BBrowser can assist the users to select any group of cells of interest to perform downstream analyses.

**Figure 6.**
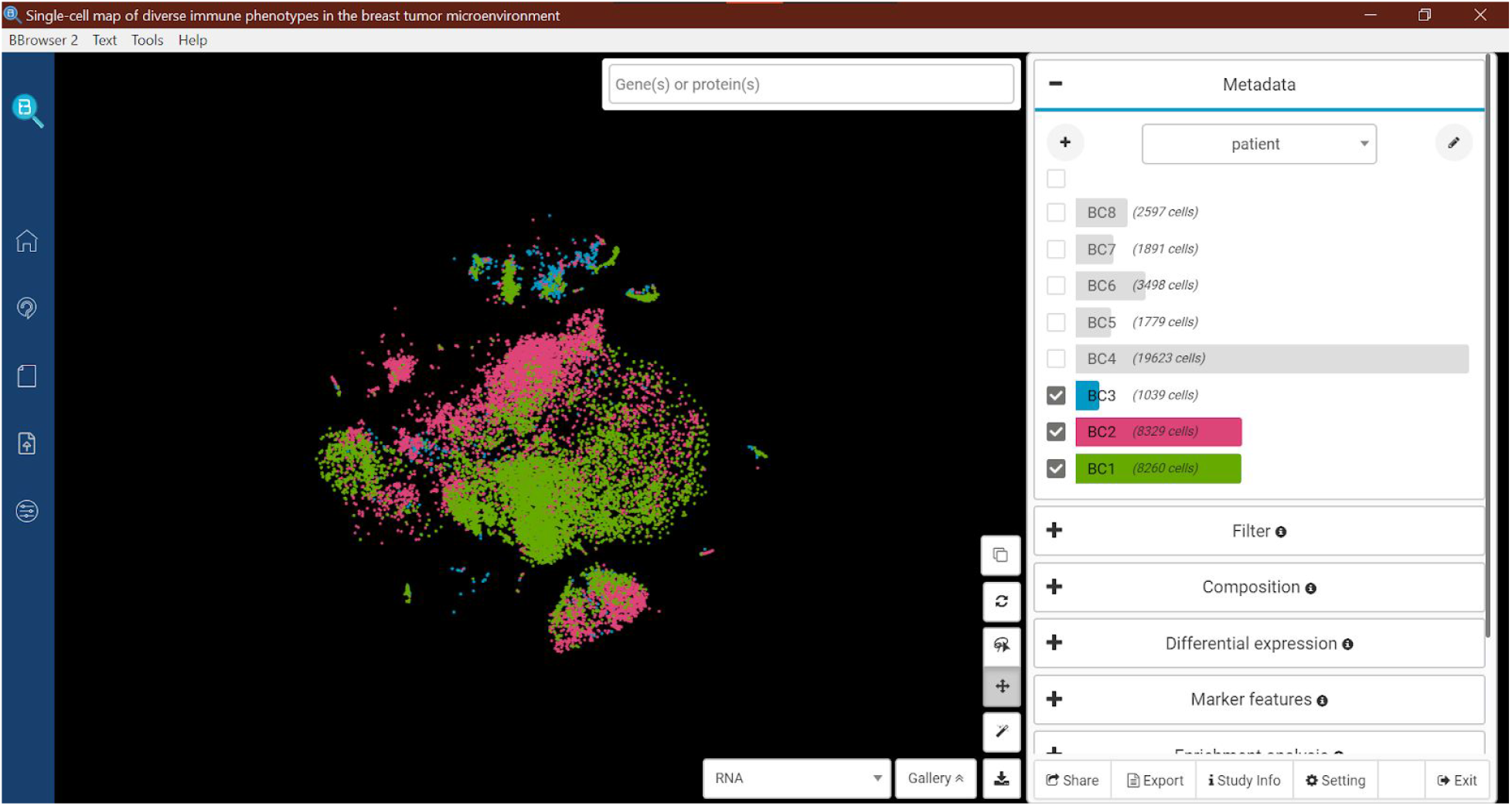
A basic cell filtering the Main Plot using the Metadata panel. The t-SNE is now colored by patient ID and only shows the cells from patient BC1-3.

**Figure 7.**
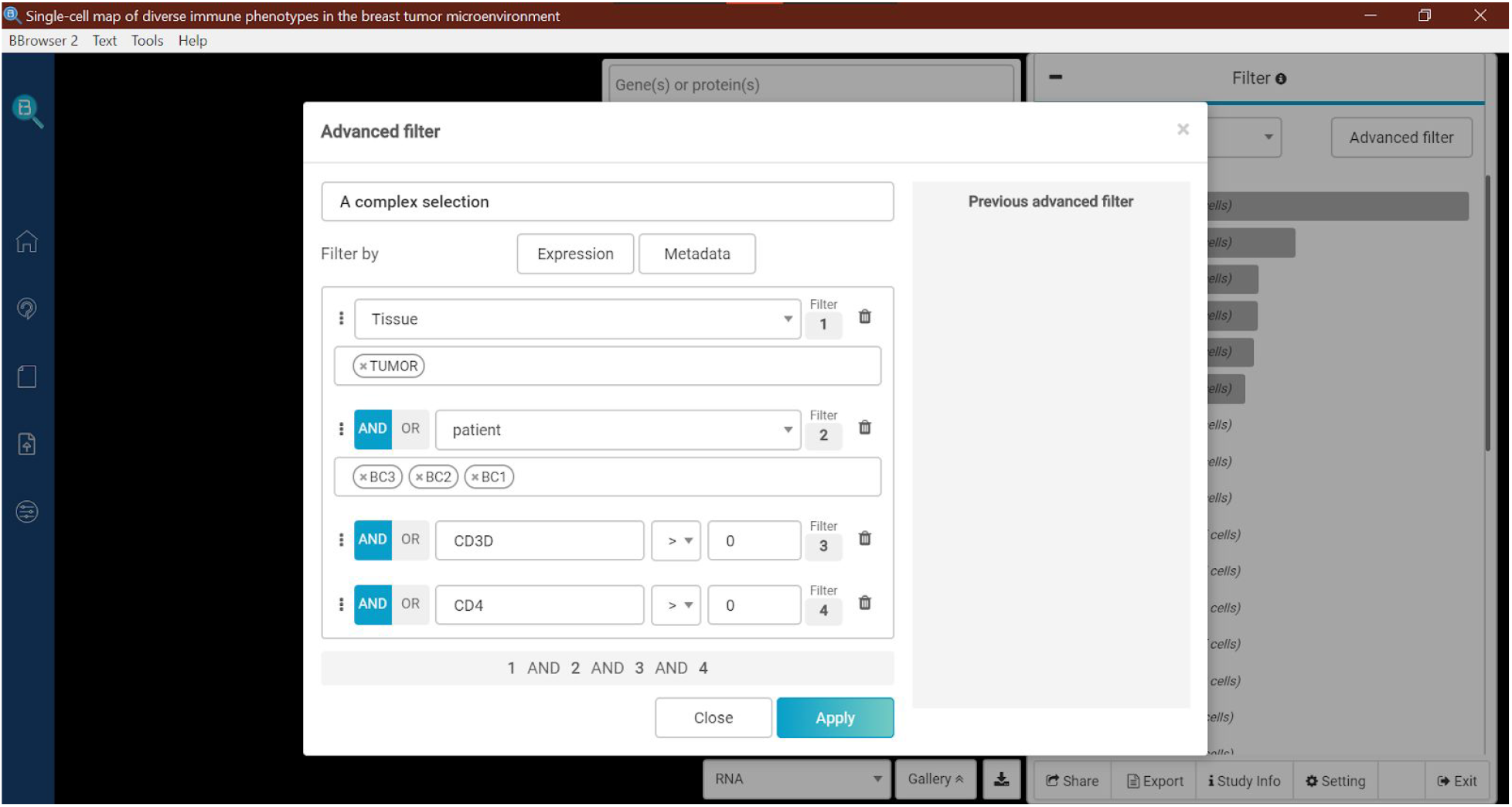
An advanced cell filtering in the Main Plot using the Filter panel. The interface shows a combination of a complex selection: only CD3D+ CD4+ cells from the tumor samples of patient BC1-3.

### Common Analyses

Given a selection of cells, the user can do many things in BBrowser. The basic function is to create a label. This new label appears in the Metadata panel and automatically synchronizes with other panels such as the Filter panel mentioned earlier (Figure 8a). Another useful analysis is to find marker genes of the selected cells (Figure 8b). This basically means to run a differential expression analysis between the selected group and the rest of the data. By default, BBrowser uses Venice (Vuong et al., 2020), but users can choose to use different methods, such as t-test, Wilcoxon, Likelihood-ratio-test, etc. To create a more specific comparison, the user can utilize the interface of the DE panel (Figure 8c) by manually defining two groups. The list of differentially expressed genes can be exported or become the input of the Enrichment panel (Figure 8d). This panel estimates the enrichment score using GSEA (Subramanian et al., 2005) on the common genesets. BBrowser supports 2 types of genesets: the Gene Ontology (Gene Ontology Consortium, 2004) and the Reactome Database (Croft et al., 2010). The software can also include 2 layers of metadata to study changes in composition. This analysis can be done in the Composition panel (Figure 8e), in which the user can choose a second metadata to observe its composition given a selection of cells on the Main Plot. This panel also supports a shortcut to perform a differential expression analysis between any to groups in the stacked bar plot.

**Figure 8.**
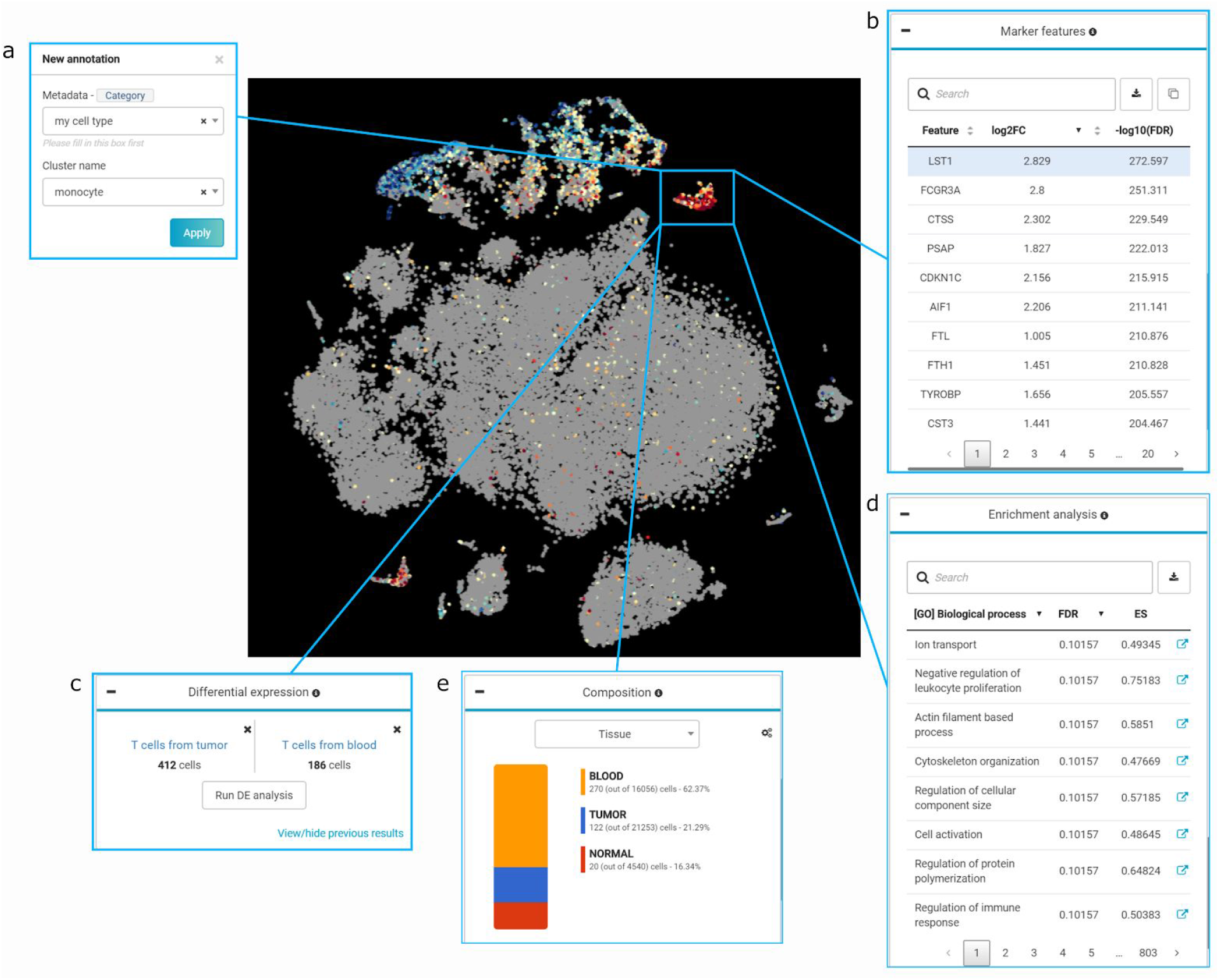
Basic analyses can be done with one click in BBrowser given a selection of cells. The center is the Main Plot where selected cells are highlighted with a cyan box. Cells are colored by the expression of LST1. Other boxes indicate the analysis panels. (a) Annotation box can create a new label to your selection. The new label will be automatically synchronized with other panels for further analysis. (b) Marker features panel can run a differential expression analysis between the selected group and the rest of the data. It creates a marker gene table that can interact with the Main Plot using genes. (c) Differential expression panel is for creating a comparison with manual selection of two groups. (d) Enrichment analysis panel can run GSEA using the output of Marker features panel. It has a list of genesets from the Gene Ontology and Reactome Database. (e) Composition panel shows the fractions of a selected metadata. This panel changes according to what cells are selected in the Main Plot.

### DE Dashboard

If the user needs to have an in-depth look into a differential expression analysis, BBrowser can further explore the result in the DE Dashboard. This dashboard enhances the focus on the two selected groups (Figure 9). The user can select a gene from an interactive volcano plot to visualize its expression via a violin or boxplot. There are also two tables for gene and enrichment analysis results similar to the normal dashboard.

**Figure 9.**
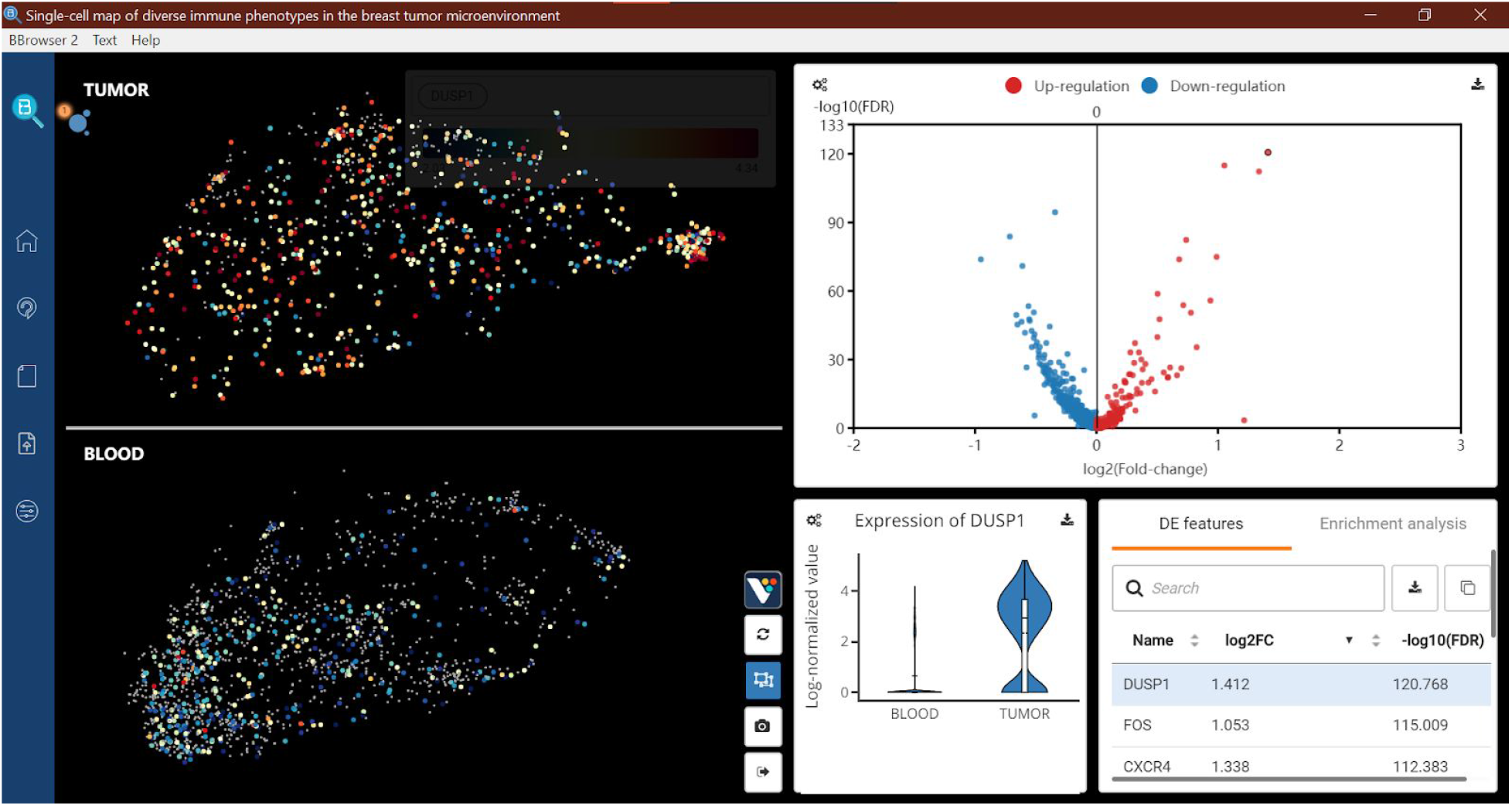
The interface of DE Dashboard in BBrowser. This dashboard enables a focus view on the comparison of two groups. It has an interactive volcano plot, where each data point is a gene in the table beneath. Clicking on any gene in this volcano plot will display the expression of that gene in the main scatter plot and the violin plot.

### Heatmaps

Besides coloring the expression of genes on the scatter plot, BBrowser supports heatmaps to better reveal expression patterns across different conditions using the Marker panel. By selecting a target metadata, for example, the graph-based clustering results, the Marker panel will create a collection of marker genes for all clusters and store them in the Gallery. The user can then add all the genes in this collection to the query and create a heatmap (Figure 10). This heatmap can be the expression values or the z-scores across cells. To better reveal the expression patterns, the user can sort the heatmap using hierarchical clustering methods.

**Figure 10.**
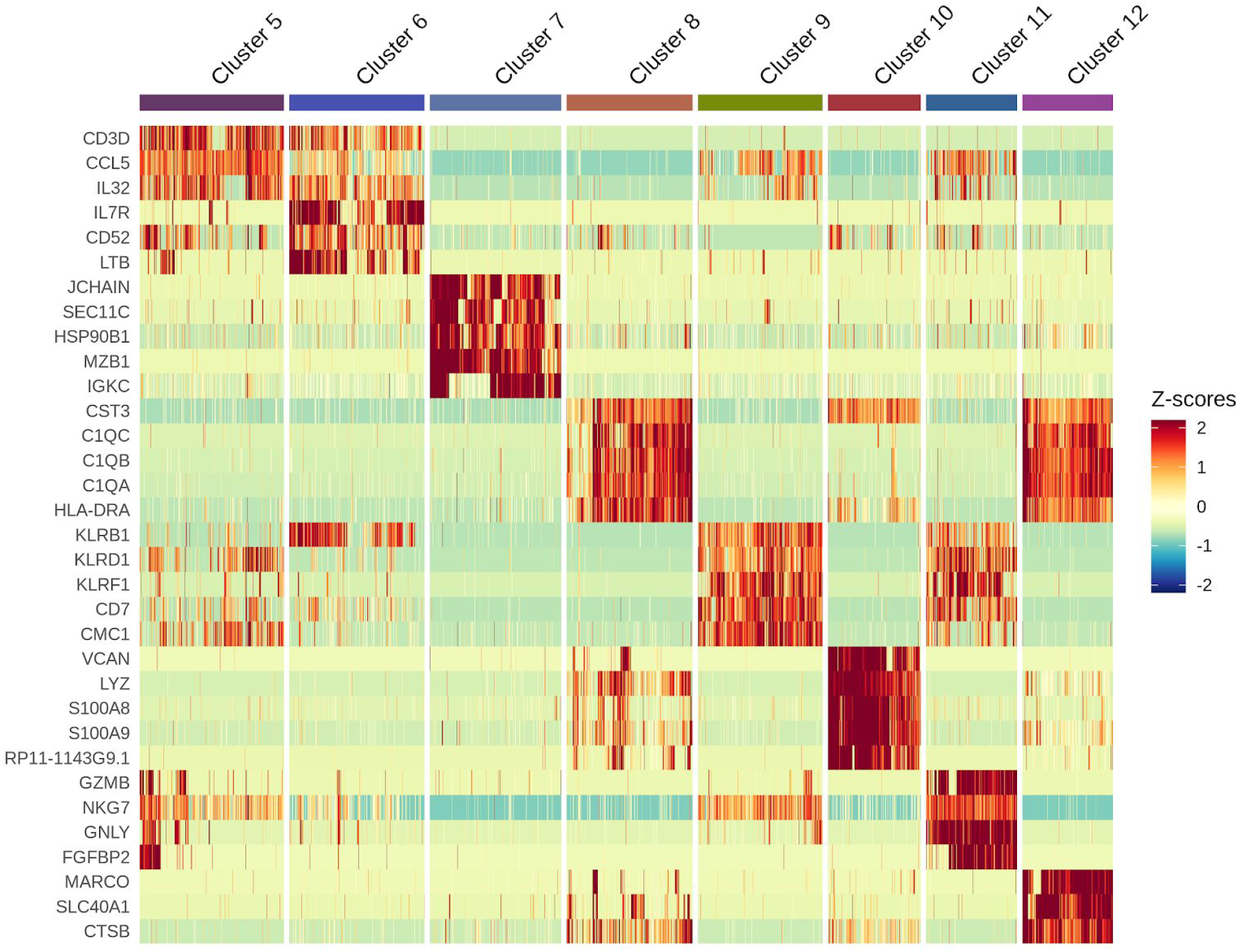
A heatmap in BBrowser using a subset of clustering results (cluster 5 to 12) with the marker genes for each cluster. Rows and columns are genes and cells respectively. Color indicates values of z-score.

### Multiple Genes

Heatmap is not the only way to analyze multiple genes in BBrowser. If the user provides two genes in the query, the software will show the color scale using the log2 of ratio. This metric helps to detect dual expressors (Figure 11a). If the user provides more than two genes, the software will show color scale as the sum of expression normalized by the total count (Pont et al., 2019). This signature score indicates the cells that are more likely to express the genes in the query (Figure 11b). Another interesting analysis for multiple genes is the gene-gene correlation matrix. The user can create a series of correlation plots from the current metadata. Users can choose different correlation metrics, including Spearman, Pearson, Kendall tau, or Jaccard Index (Figure 12). These lists of genes can be stored under the Gallery so that the user can quickly repeat the previous analysis in that data set (Figure 13).

**Figure 11.**
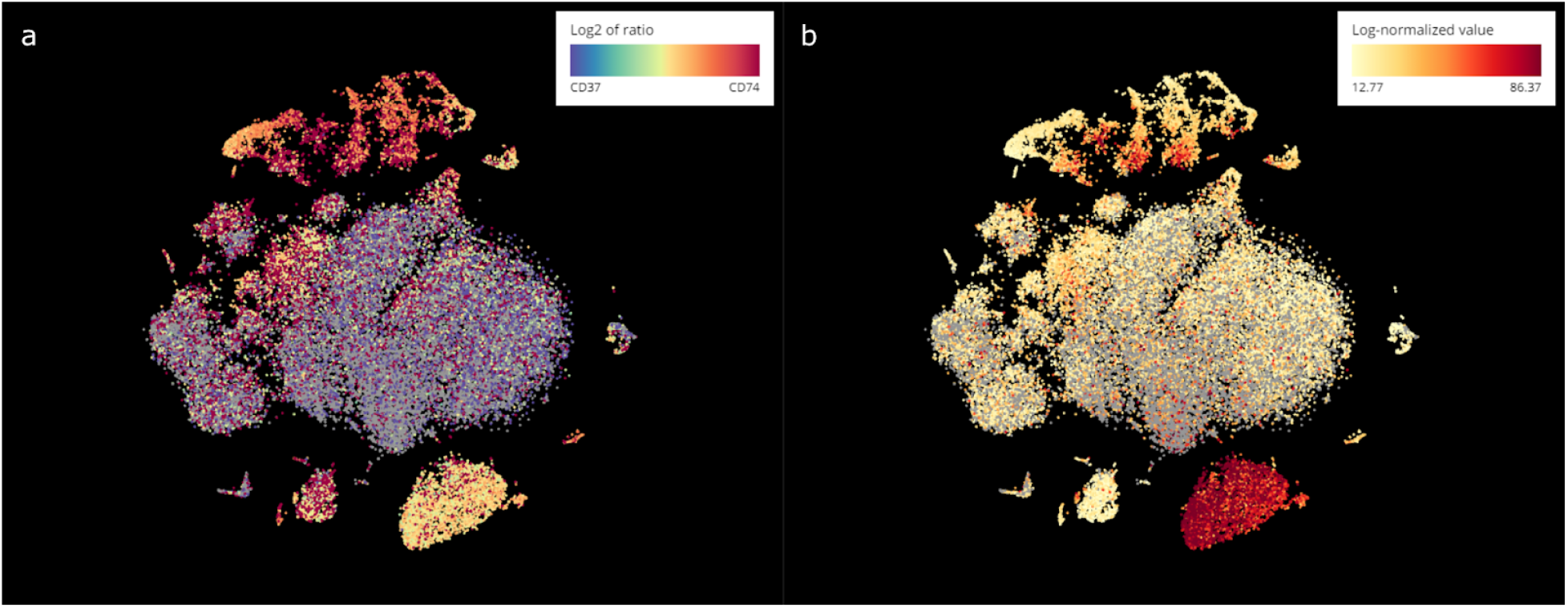
The Main Plot shows the color scale of multiple genes. (a) The color indicates the log2 ratio of expression between CD37 and CD74. Cells in gray do not express both of these two genes. (b) The color indicates the signature score from the log-normalized expression of CD37, CD74, and CD79A.

**Figure 12.**
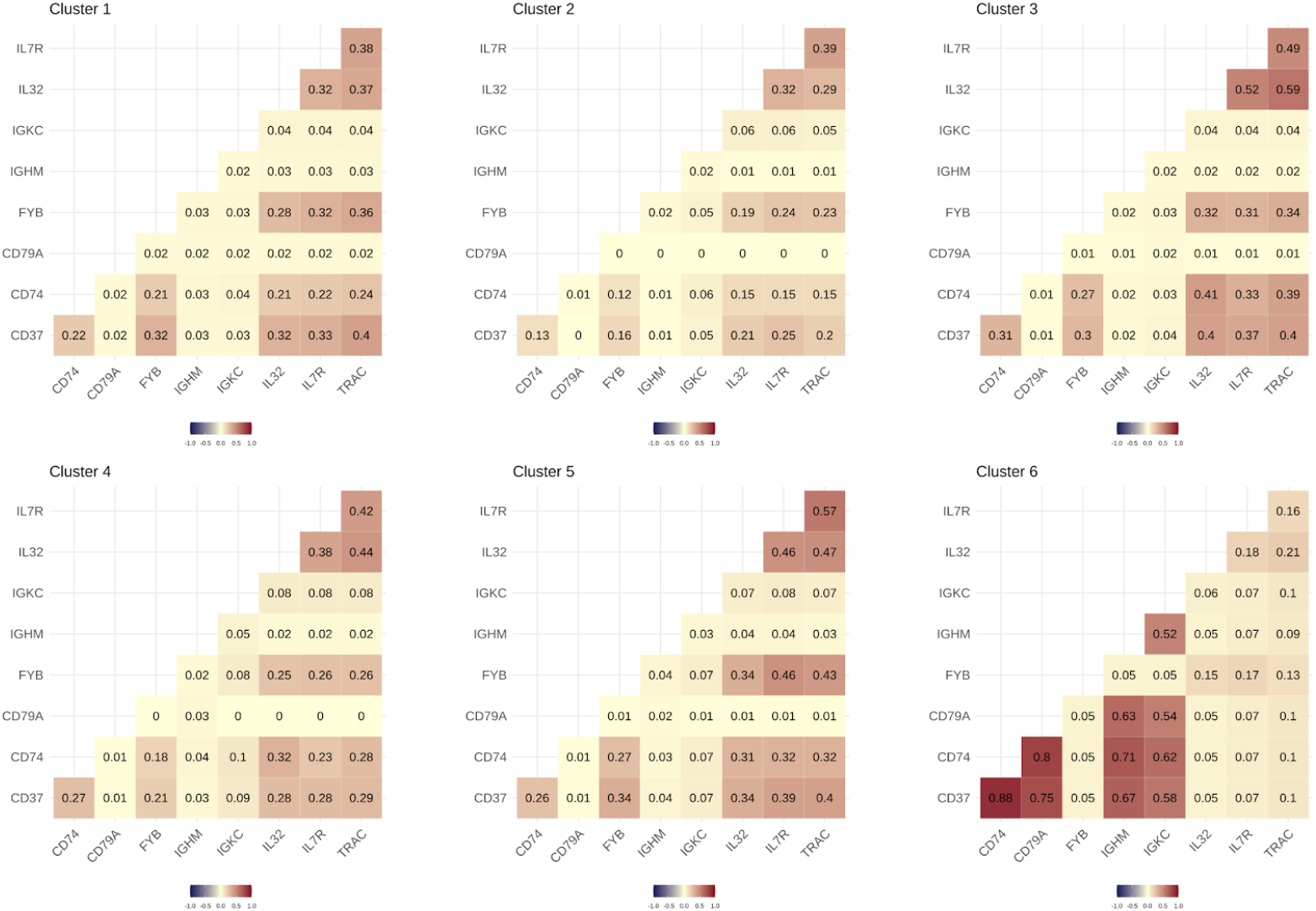
The correlation plots in BBrowser for 6 clusters using the expression of 9 genes. The color scale indicates the Jaccard index and can be changed to Spearman, Pearson, Kendall tau.

**Figure 13.**
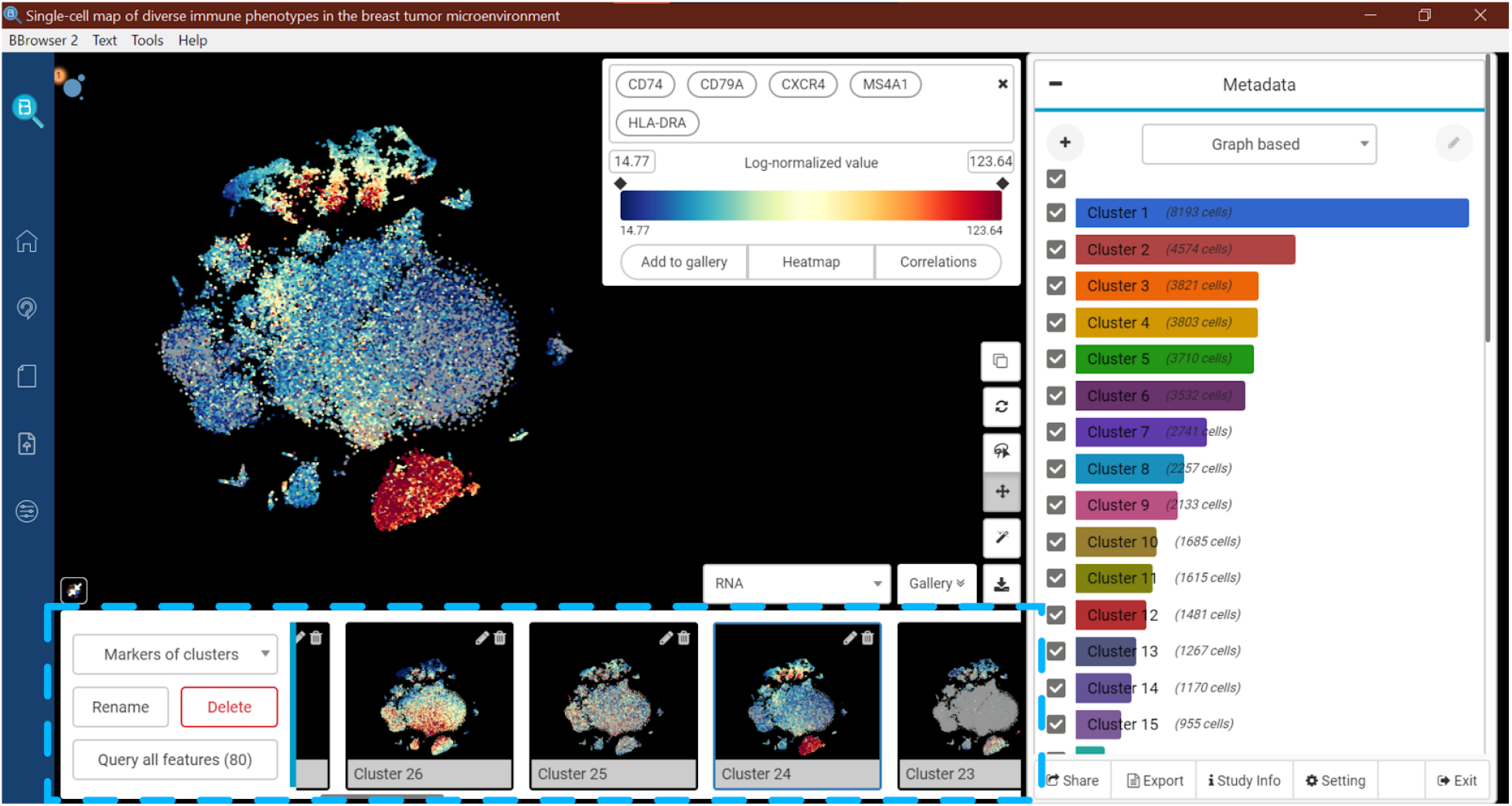
Gallery of features in BBrowser. The gallery section is highlighted with a dashed box. This is where the user can store a gene set or a list of markers for a specific data set.

### Interactive Snapshots

Another complex visualization is the ability to splitted the Main Plot and interact with all the splitted plots simultaneously. The user can apply a filter for the Main Plot, then create an Interactive Snapshot (Figure 14). This function generates a secondary window that listens to the events in Main Plot. Thus, when the user interacts with the Main Plot, such as querying a gene or analyzing a group of cells, all the snapshots will react coherently. It is particularly helpful when the user has to deal with multiple types of omics at once, such as in the spatial transcriptomics data. An interactive snapshot can simultaneously visualize a t-SNE plot while the user interacts with the spatial coordinates (Figure 15).

**Figure 14.**
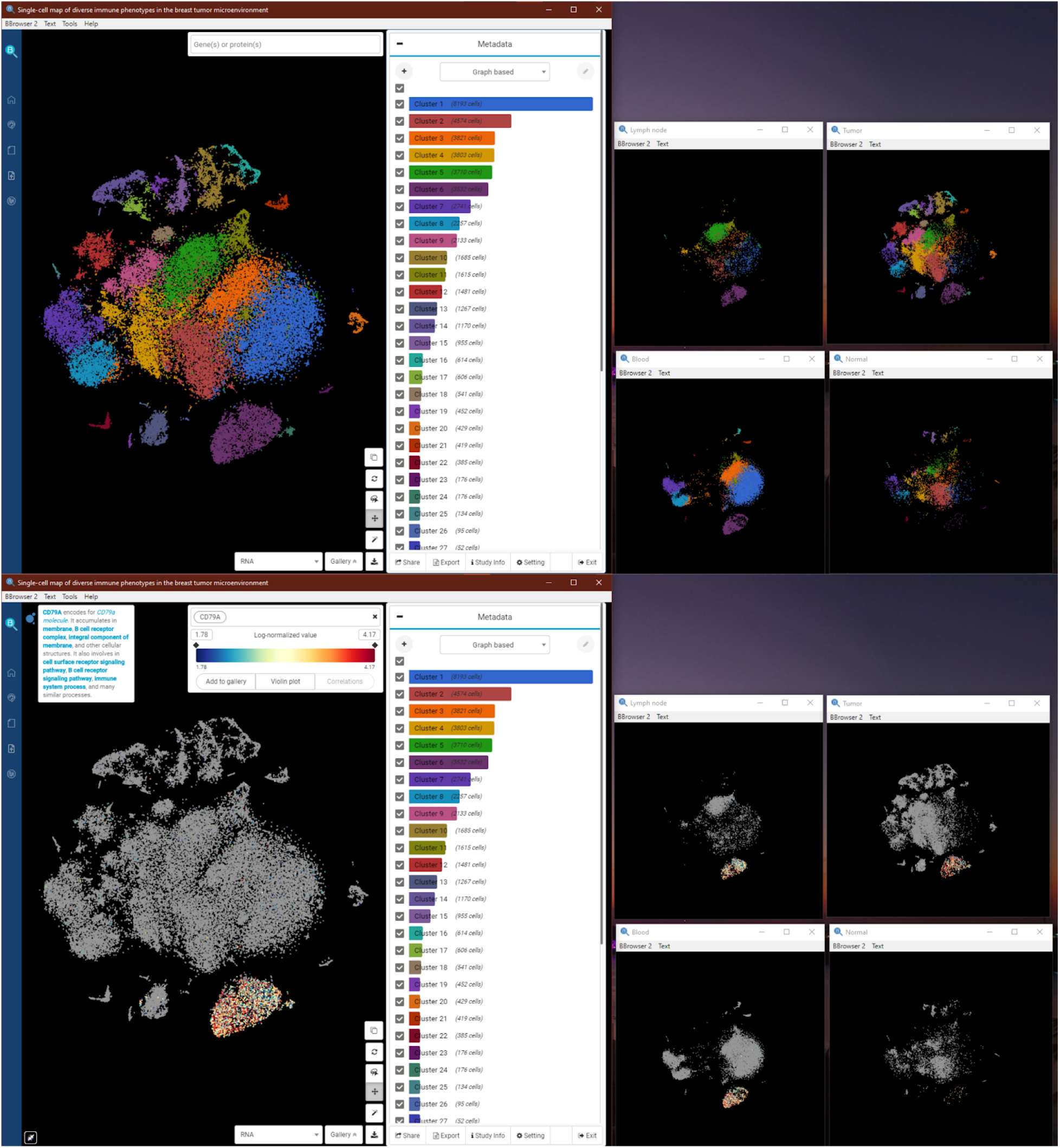
Two screenshots of the monitor to illustrate the usage of interactive snapshot. The upper screenshot shows a primary window (left) with 4 interactive snapshots from 4 different tissues (Lymph node, Tumor, Blood, and Normal). The lower screenshot shows the behaviours of these snapshots when the user changes the color of the Main Plot into the expression of CD74A. All snapshots change color accordingly. This behavior can also apply for the change of color in selection, metadata, composition, and clonotypes.

**Figure 15.**
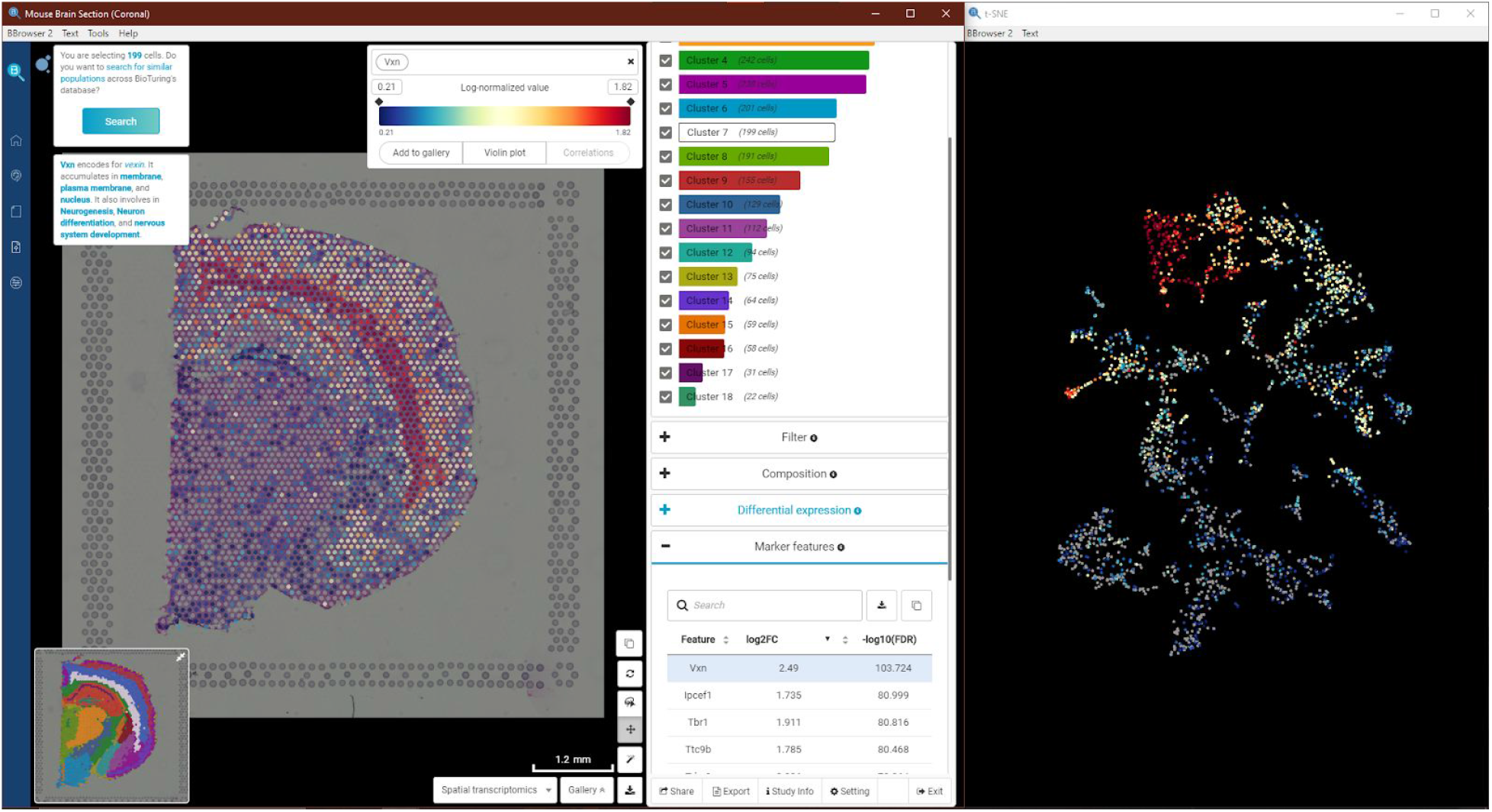
Illustration of interactive snapshots in BBrowser for spatial data. A snapshot was created for the t-SNE of the data spot (right). This snapshot shows the same gene, Vxn, as the primary window. The <u>data</u> was generated by 10X Genomics.

### Clonotypes

For data sets with TCR or BCR sequencing data, the user can see the clonotype table using the Clonotype panel (Figure 16). This panel consists of an interactive table that changes accordingly with the Main Plot. It has a mini composition chart using the metadata that can be chosen in the Metadata panel. The user can count the clonotype using the identity of both or one type of chain in the receptor. We incorporated a VDJ database (Shugay et al., 2018) inside BBrowser so that it can tell the relevant MHC molecules and antigens given a CDR3 sequence.

**Figure 16.**
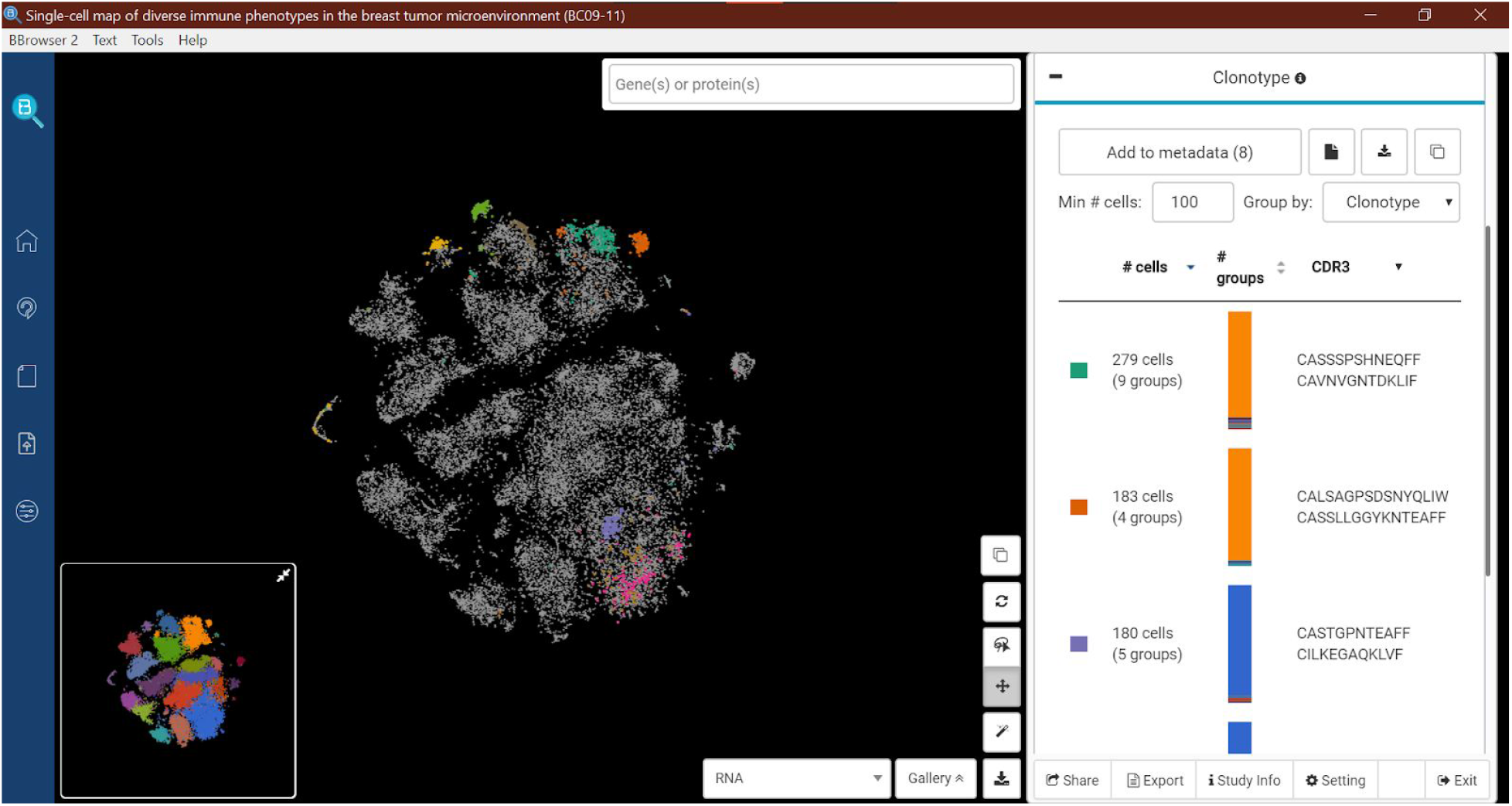
The interface of BBrowser with the Clonotype panel opened. This panel colors the Main Plot with the clonotype ID. It also shows the clonotypes in a table with details about CDR3 as well as the relevant antigen information. A mini composition plot is included to estimate the fractions of current metadata.

### Subclusters

To assist performing studies on sub populations of cells, users can perform sub-clustering in BBrowser. Once a subcluster is created, it will be a part of the original data set so that their metadata will synchronize with each other. A subcluster inherits all the functionality of the original data set, thus, enables a wide range of applications to explore the heterogeneity of a single-cell data.

## PROCESS IN-HOUSE DATA

In addition to accessing the public database, BBrowser can host and analyze private single-cell data sets. The software supports various types of input data at different stages (Figure 17). The user can submit raw sequencing data in FASTQ format, as well as processed files. Raw data are aligned and quantified by Hera-T, an accurate and efficient algorithm for single-cell sequencing data (Tran et al., 2019). Already-processed data (i.e., expression matrices in MTX, CSV or TSV) can also be loaded into the software. Data with antibody-derived tags such as CITE-Seq (Stoeckius et al., 2018) or REAP-Seq (Peterson et al., 2017) will also be loaded from MTX format. For spatial data, BBrowser has a separate interface to load spatial data, and aims to support 10X Genomics Visium platform and Nanostring GeoMx DSP.

**Figure 17.**
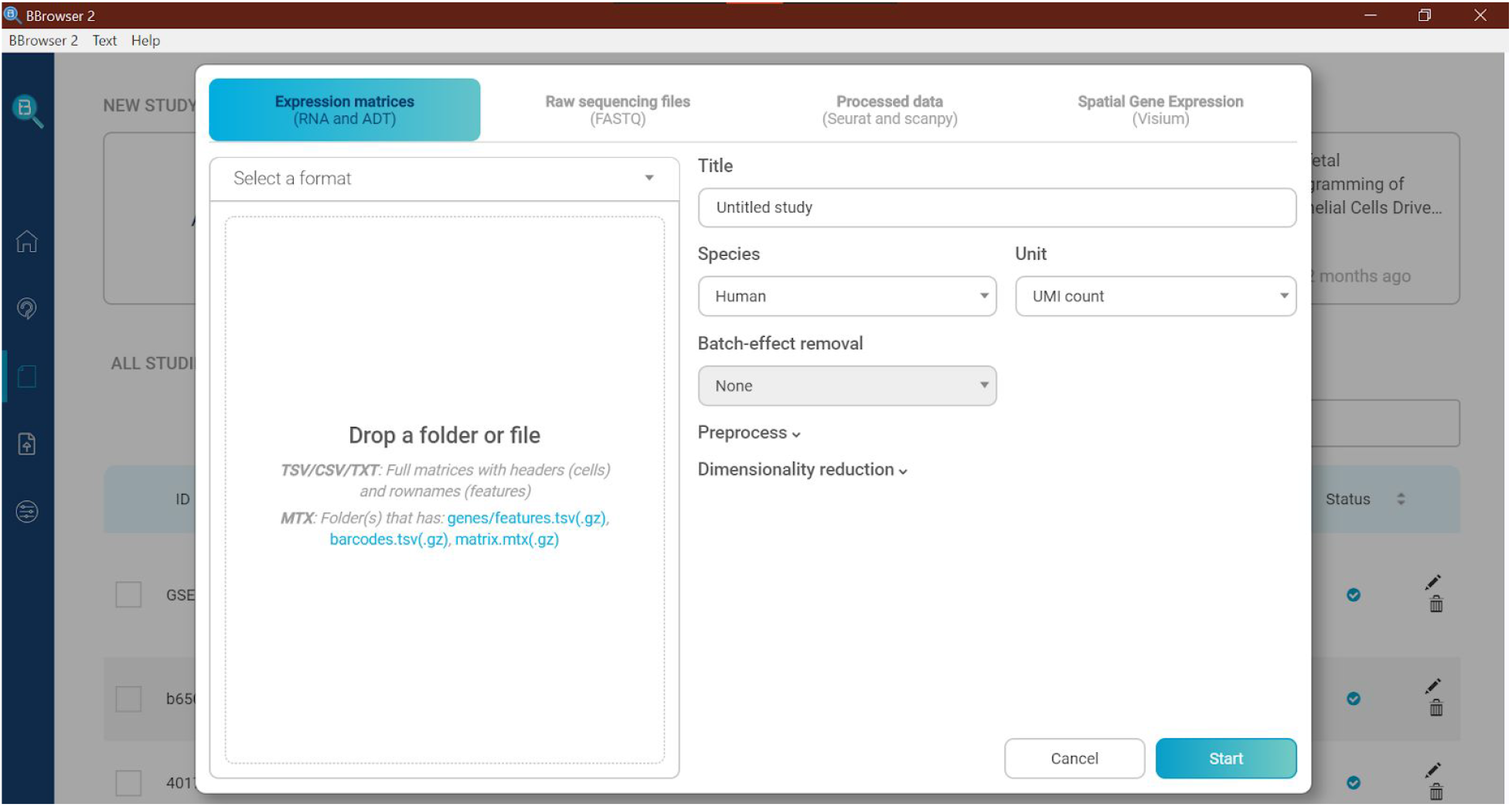
The interface to load new data into BBrowser. This interface supports input from raw sequencing files (FASTQ), expression matrices (MTX, text-delimited), and processed data (RDS and H5AD). There is a separate section for spatial transcriptomics where the user can load its spatial images.

The user can further customize additional parameters to process the data. There are basic parameters for filtering such as the number of expressed cells or genes, total percentage of mitochondrial expression, or number of highly variable genes. More advanced integration of multiple batches can be done with CCA (Stuart et al., 2019), MNN (Haghverdi et al., 2018), or Harmony (Korsunsky et al., 2019). For dimensionality reduction, BBrowser can perform t-SNE (Maaten et al., 2008) or UMAP (McInnes et al., 2018) with custom perplexity. All stochastic processes in BBrowser have a modifiable random seed to ensure reproducibility.

However, we acknowledge that there are many different combinations of methods to process a single-cell data set. To allow this flexibility, BBrowser supports importing data processed by common single-cell packages such as Seurat (Stuart et al., 2019) or scanpy (Wolf et al., 2018). The user can process data with their favorite pipeline and store results in standard formats such as RDS (in R) or H5AD (in python). BBrowser will extract embeddings, metadata, and normalized values and use them for downstream visualizations.

## INTEGRATION

In its initial design, BBrowser was a desktop application with data storage and analysis performed on local computers. This model limits the ability to analyze big data sets and to share data. To address this limitation, we create a separated layer for computational and data storage. BBrowser communicates with this layer via API calls. Hence, the computational and data storage can be anywhere, either the local computer or a remote server. This architecture is called *BioTuring Enterprise Server Platform* (BESP) (Figure 18). This solution helps users from a laboratory, an institute, or a private company to process large single-cell data concurrently and centralize those data on their high-performance server.

**Figure 18.**
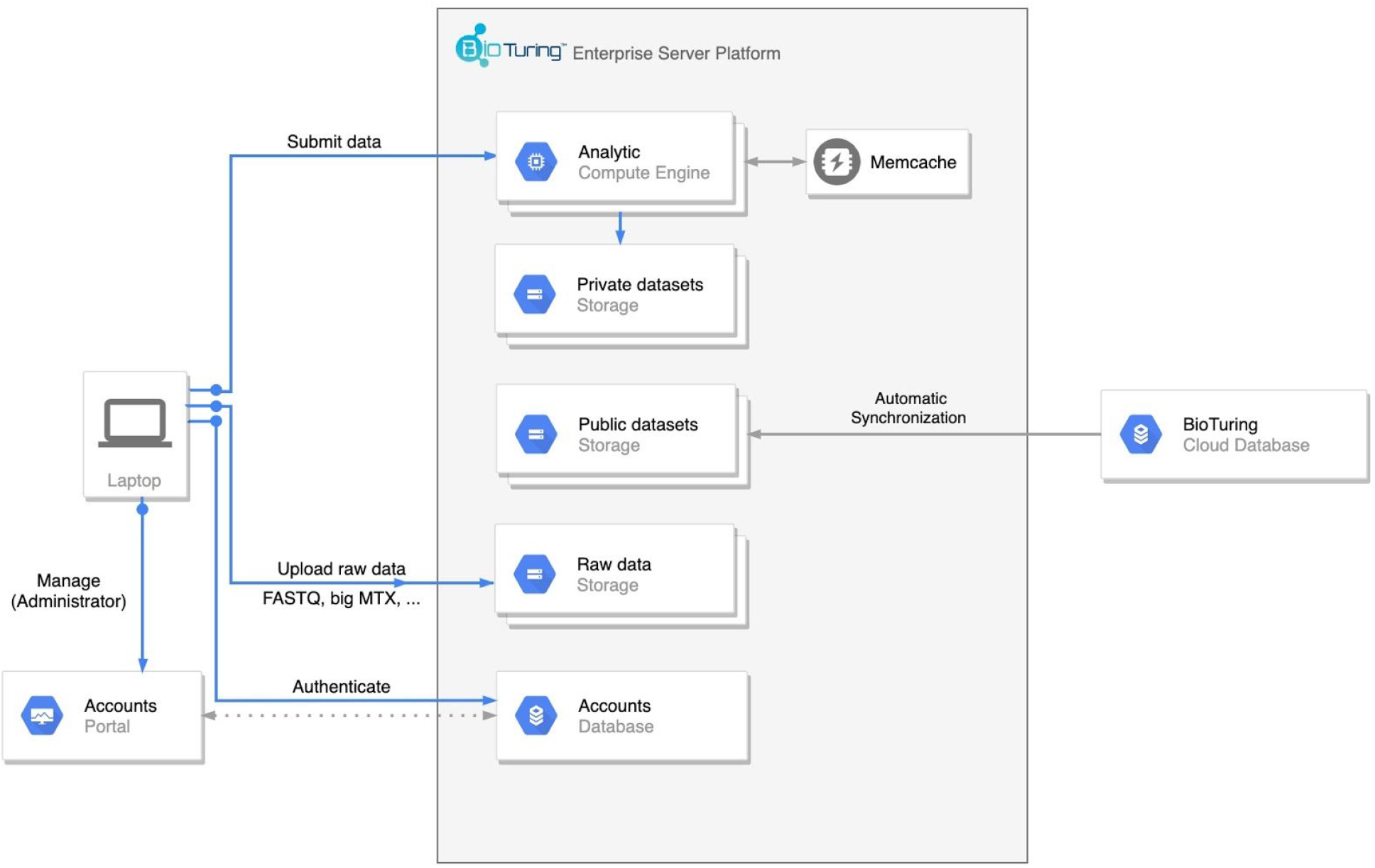
Architecture of the BioTuring Enterprise Server Platform (BESP). Each desktop version of BBrowser (laptop) is a client that requests BESP for data processing and storage. The clients are managed by the *Accounts Portal*, where the administrator can set read/write permission, create groups, update, or reconfigure BESP.

By centralizing data on BESP, users can share data with other members in their groups. For each data set, different users can use BBrowser to create multiple analyzes and therefore, change the original data. As the size of each single-cell data set is big, replicating a copy of the data for each shared user is expensive in both disk space and copy time. To address this problem, we implemented a fast and resource-efficient sharing method by separating static and versatile files.

Allowing users to share datasets with each other enhances communication but also raises a problem about data reproducibility. To address this problem, we developed a version control system on the BESP server. Similar to git, the history of each dataset is a directed graph of snapshots (Figure 19). Each node on the graph is a shared snapshot of one member in the group. A directed edge *u->v* on the graph represents that the snapshot *u* is the original data of *v*. Users are able to download a particular snapshot on the graph, then can make some changes and share it back to the server. By doing that, they created a new branch on the graph, and thus expanded a new path of development for the dataset.

**Figure 19.**
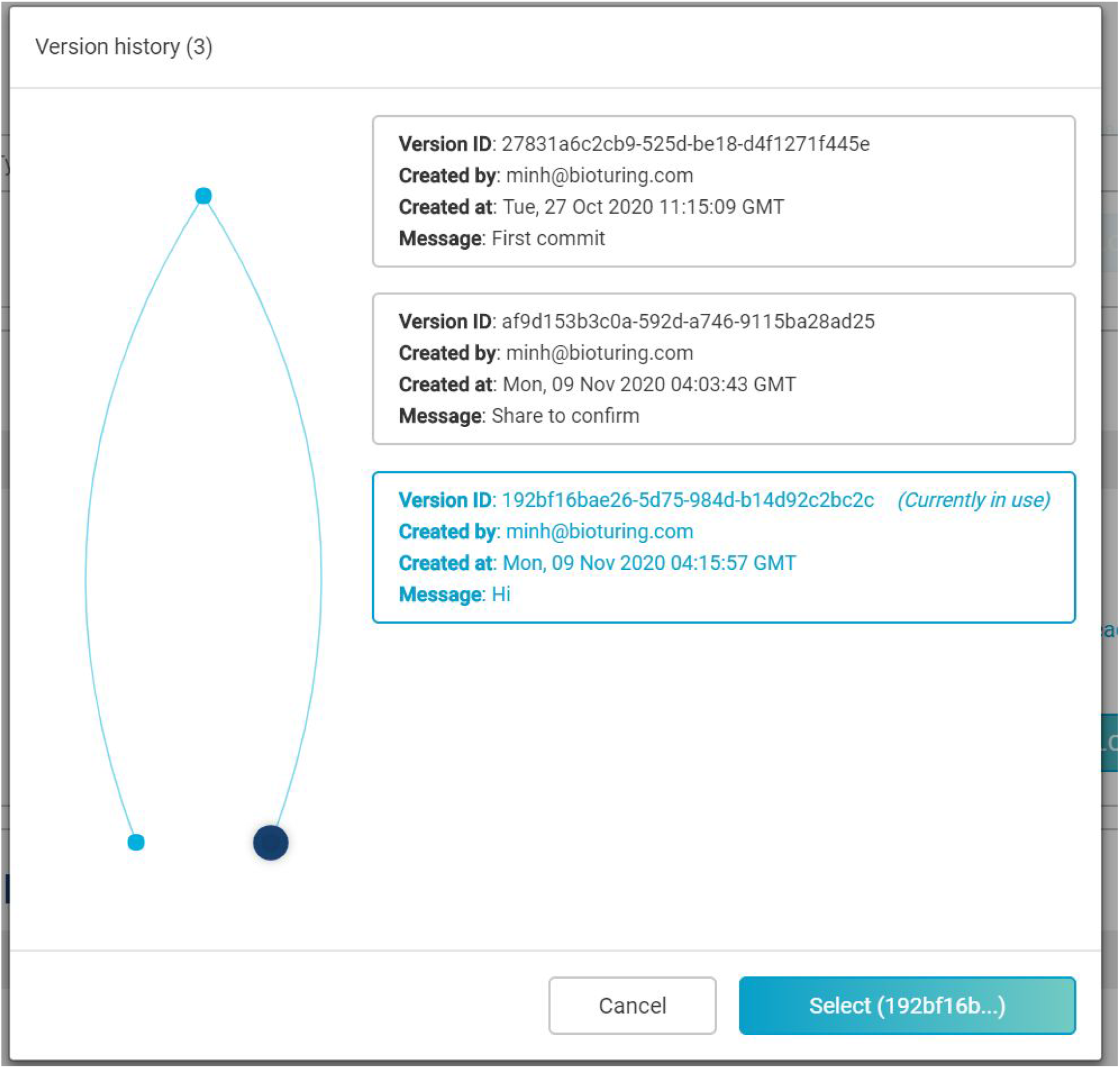
A visualization of a history of a shared data set. A graph (left) shows each version (node) connected to its respective origin. A list of versions (right) shows general information. The user can go back to any previous version of the data set.

To allow users to utilize the power of cloud computing to process a large number of data sets by customizable pipelines, we designed a Batch-Submission framework that utilizes Cromwell (Voss et al., 2017) as the backend. Cromwell framework consumes workflows written in WDL, or CWL. Those languages are easy to understand and are popular in large-scale data processing. Cromwell is being used in Cumulus (Li et al., 2020) -- a multi-purpose and cost-effective single-cell analysis workflow to process data for the Immune Cell Atlas project, which involves the transcriptomic profile of more than 1.7 million cells.

The separation of the computing and data layer to server mode allows us to integrate multiple services on private data sets. For instance, *Talk2Data* and *Cell Search* can be deployed on BESP to index and return results using proprietary single-cell data of the institutions/companies. In particular, users can build custom indexes for *Cell Search* and *Talk2Data* to analyze/visualize on their own data and cell annotations.

## CASE STUDIES

We illustrate the usage of BBrowser to reproduce and analyze deeper into a single-cell dataset.

### Case 1: Characterize breast tumor microenvironment

Single-cell technologyprovides unprecedented resolution to study the tumour microenvironment. In 2018, Azizi and colleagues applied this technique to obtain a large-scale characterization of breast tumor immunology (Azizi et al., 2018). The data set that we are going to use from this work will be the samples from patients BC01 to BC08, comprising 47016 cells in 4 different tissues. In BBrowser, this data set has an ID “GSE114725”.

Let us imagine that there is no cell type label in this data set. This is the common scenario when the user first loads a single-cell data into a software. Cell typing can be done numerous ways with BBrowser. The first way is to query for a known marker gene, for example CD79A for B cells. BBrowser can show the expression value on the t-SNE, as well as a violin plot across the graph-based clusters (Figure 20). The second way is to utilize the *Marker* panel to look for the marker genes of a particular cluster of cells. For example, BBrowser found cluster 23 highly expresses IRF8, IRF7, LIRA4, and GZMB, which indicates it might be a cluster of plasmacytoid dendritic cells. The third way is to use Cell Search, a function we mentioned earlier, can also assist the cell typing process. We used this function on cluster 21 and found 107,698 similar cells in 11 different studies, all labeled with “monocyte”.

**Figure 20.**
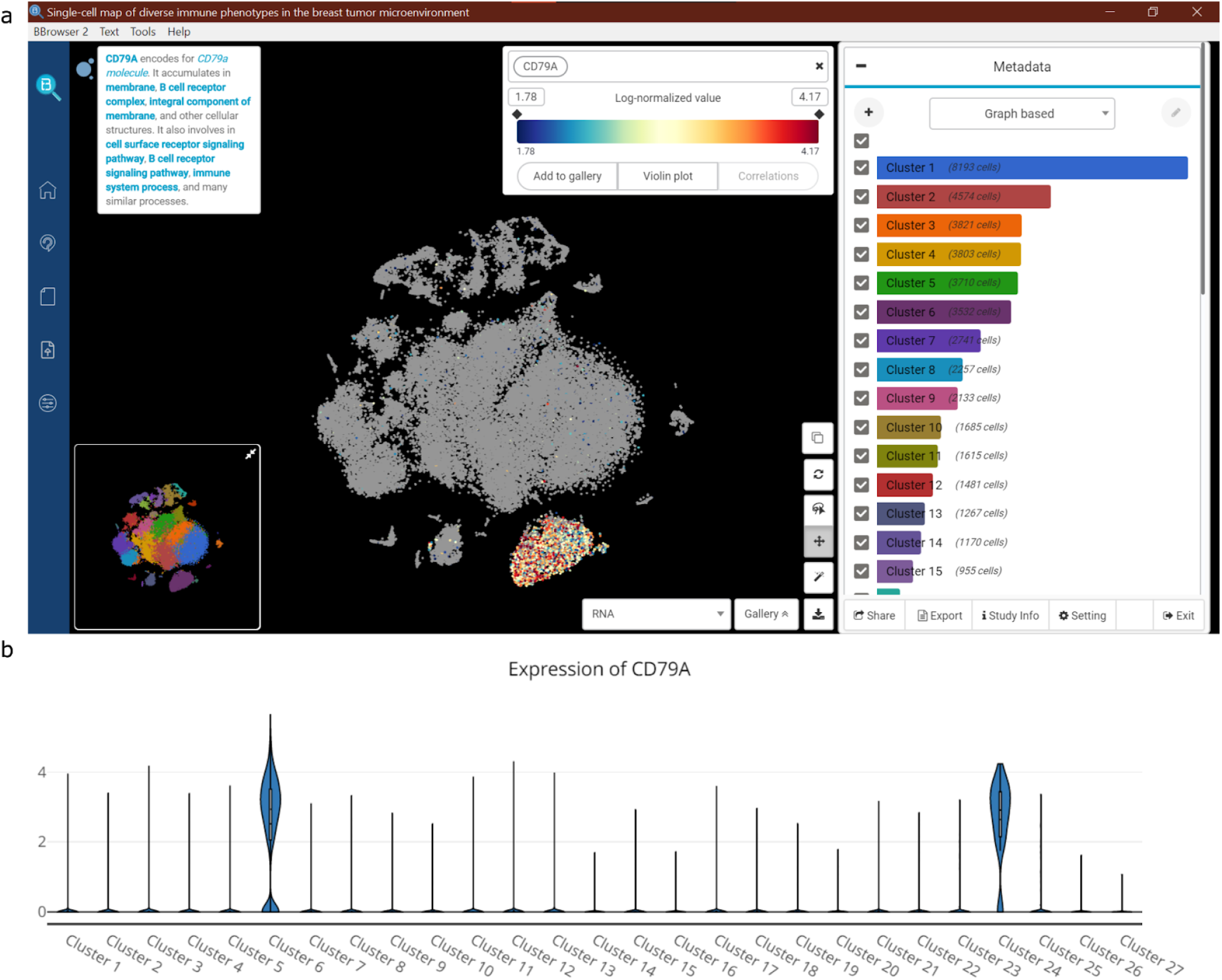
B-cell identification using a known marker gene, CD79A, in BBrowser. The software shows the expression value on the t-SNE, as well as a violin across all graph-based clusters.

After having all the cell types identified, we also re-explored the tumor microenvironment. We found that each tissue (except normal tissue) has a unique prevalent cell type. For example, B cell is more abundant in lymph nodes, while the cytotoxic T cell is more abundant in tumors (Figure 21). These findings are consistent with the paper. The paper also mentioned that there is a specific naive CD4+ T cell in blood compared to the ones in lymph nodes. We also observed two separate clusters of naive CD4+ T cells and even went further using the DE dashboard of BBrowser. We detected one marker gene, CXCR4, which only up-regulates the naive CD4+ T cells from lymph nodes. This is consistent with an earlier research that CXCR4 plays an important role for T cells to enter afferent lymphatics (Geherin et al., 2014).

**Figure 21.**
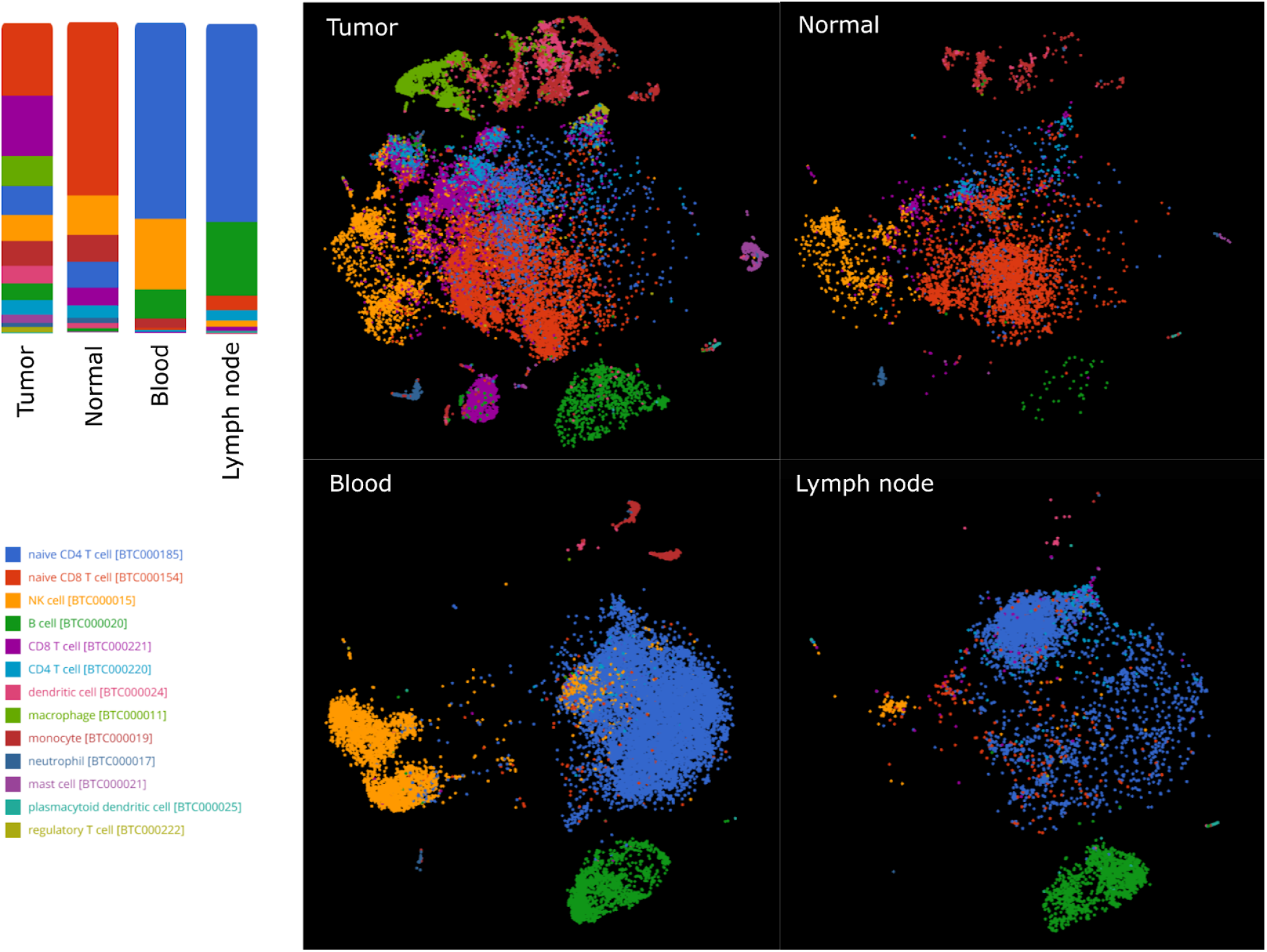
An illustration of visualizations generated by BBrowser to study the cell type composition of different tissue. Top left: a composition of cell types in 4 different tissues. Right: The t-SNE plot of 4 different tissue colored by the cell type annotations. Bottom left: the color legend.

### Case 2: Discover dual expressors T- and B-cell receptors hidden in published datasets

In every biology textbook, expression of the B cell receptor (BCR) defines B cells, and the T cell receptor (TCR) defines T cells. However, in 2019, Ahmed and colleagues discovered a strange cell population in Type I Diabetes that coexpresses the BCR and TCR, and key lineage markers of both B and T cells (Ahmed et al., 2019). In this case study, we compared the T cell populations in lung tumors vs normal lung tissue from all BioTuring lung cancer single cell data and surprisingly found that B Cell Receptor (BCR) light chain and heavy chain genes (specifically, IGKC and IGHA1) to be up-regulated in lung adenocarcinoma.

We compared the T cells from the tumor and normal lung tissues of a data set in a study of lung tumor and brain metastasis (Kim et al., 2020). We found significant up-regulation of immunoglobulin genes in tumor-infiltrating T cells. These most significant genes included IGKC, IGLC2, IGHA1 and IGHG3. Meanwhile, IGKC, the most significant gene, was also highly up-regulated in the T cells of brain metastasis, compared with those in the normal lung tissues (Figure 22). Our composition analysis for T-cells in tumors also showed that CD8+/CD4+ mixed T helper cells were the subtypes that exhibited the highest expression of IGKC (Figure 23).

**Figure 22.**
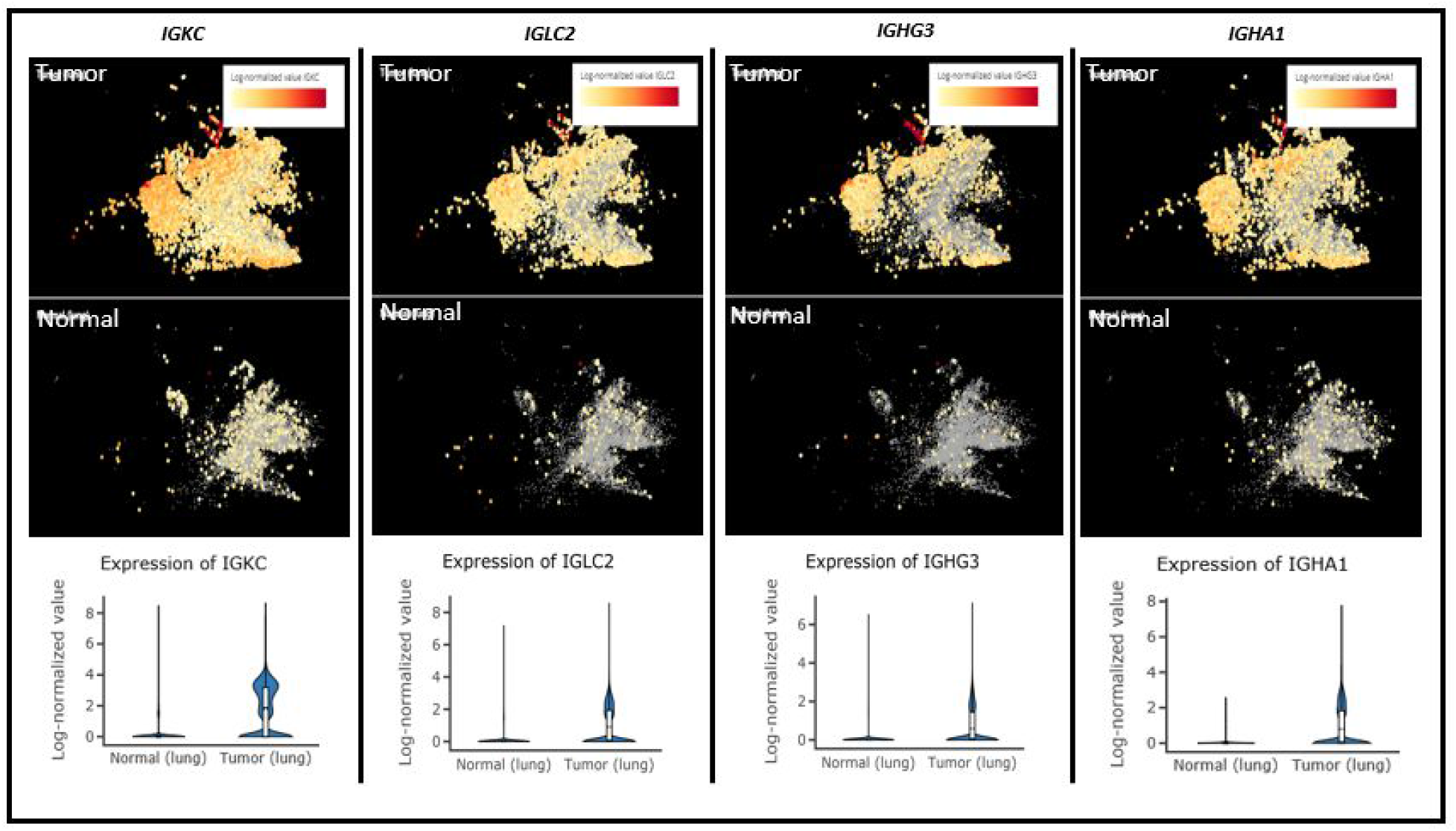
Top 4 Immunoglobulin genes up-regulated in Tumor T cells from left to right: IGKC, IGLC2, IGHG3 and IGHA1. Upper panel shows T cells from lung cancer while the lower panel shows T cells from normal lungs.

**Figure 23.**
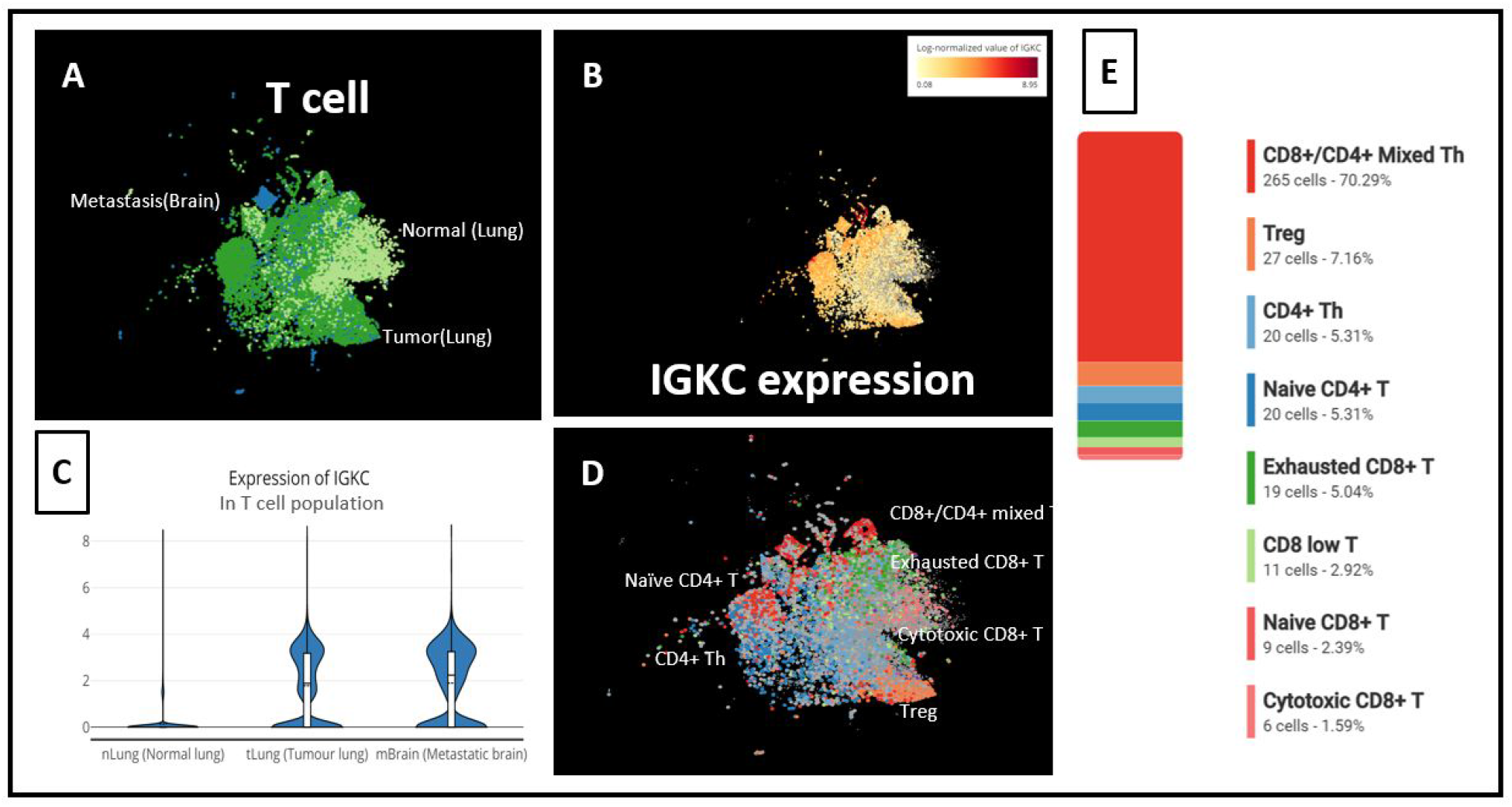
(A) t-SNE of T cells from 3 groups: Brain Metastasis, Normal (Lung) and Tumor (lung), (B) IGKC expression level in T cells, (C) IGKC expression across T cells from Brain Metastasis, Normal (Lung) and Tumor (lung), (D) T cell subtypes identified in the study, (E) Composition analysis showed that top IGKC expressors in Tumor (lung) were mainly CD8+/CD4+ mixed T helper cells.

In another data set of lung cancer metastasis (Laughney et al., 2020), we also compared CD3D+ cell populations from the tumor vs. normal lung tissue. And immunoglobulin genes are also up-regulated in the T cells from tumor and metastasis groups, compared with normal lung tissues. Interestingly, in this dataset, IGKC was gradually up-regulated in T cells from normal tissues to tumor to metastasis. In particular, this appears in the memory T cell, NK T cell, and T helper cell (Figure 24). Using BBrowser, scientists can replicate this finding without spending too much time on writing scripts.

**Figure 24.**
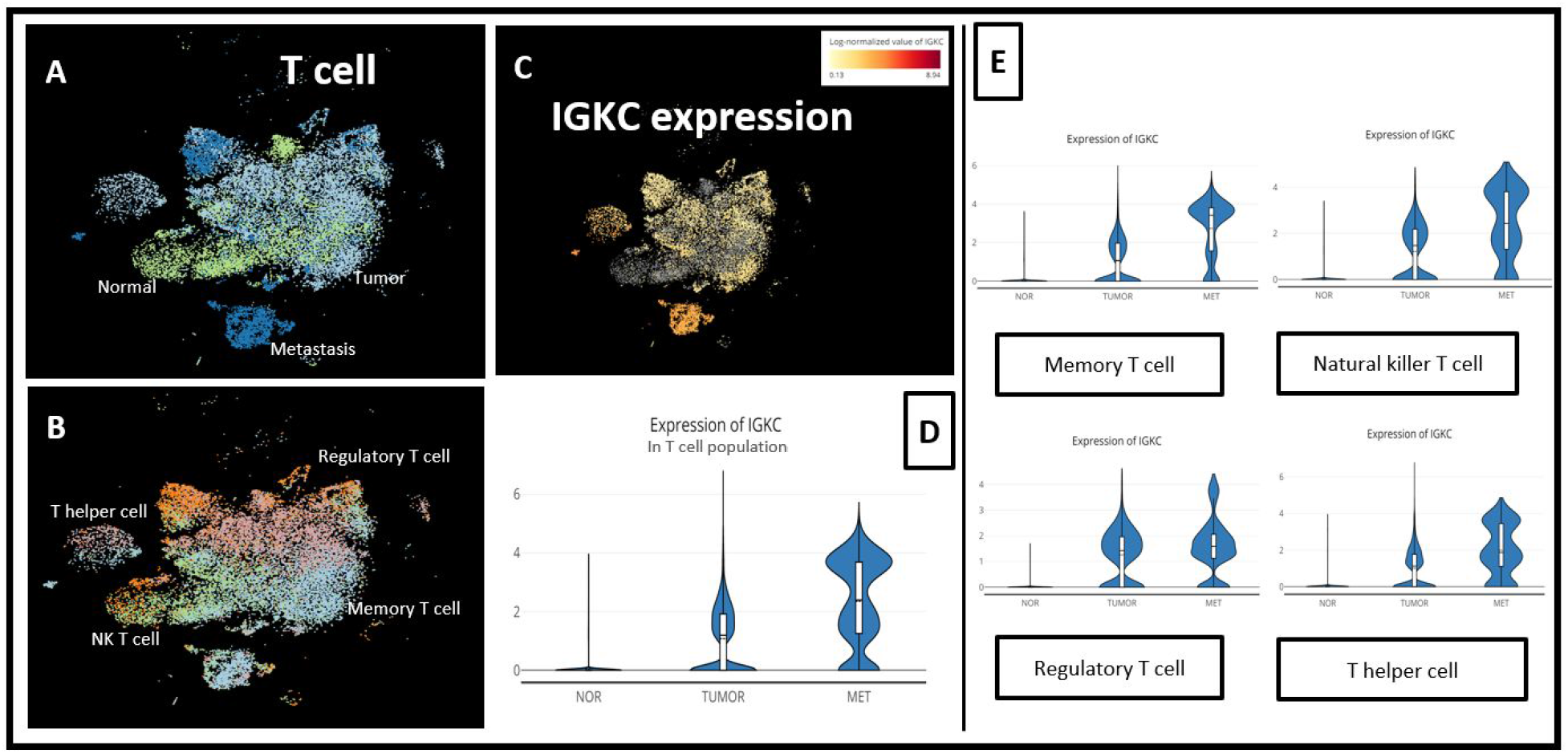
(A) t-SNE of the T cell population of 3 groups: Metastasis, Normal and Tumor, and (B) all T cell subtypes in 3 groups; (C) t-SNE of IGKC expression and (D) comparison of IGKC expression level across 3 conditions in the entire T cell population and in (E) each T cell subtype.

## CONCLUSION

BBrowser combines big computation, big data and modern visualization to help scientists interact and obtain important biological insights from the massive amounts of single-cell data. We also described the use of BBrowser to integrate with in-house pipeline, efficiently re-analyze published findings, and discover new insights.

